# An extended omnigenic model explains genome-phenome relationships for complex traits in global sorghum diversity

**DOI:** 10.1101/2024.04.29.591686

**Authors:** Zhenbin Hu, Xu Wang, Sandeep R. Marla, Jesse Poland, Geoffrey P. Morris

## Abstract

A central finding of complex trait genetics is the geometric distribution of effect sizes, but the biological basis of this phenomena is not understood. The omnigenic model (OM) could explain this architecture, with oligogenic variation arising from direct regulatory genes (core genes) and polygenic variation from indirect regulators (peripheral *trans* regulators). Plant yield is a canonical complex trait and here we tested the OM using genome-phenome analysis of biomass yield in global sorghum diversity. We used field-based phenomics to characterize dynamic growth and yield formation traits, then decomposed oligogenic and polygenic components of variation using genome-wide association studies (GWAS), genome-wide prediction (GWP), and tissue-specific transcriptome analyses. We identified major dynamic QTL, including several persistent (*Dw1*, *Dw3*) or transient (*ELF3*) QTL at known genes, consistent with the oligogenic-core of the OM. Next, we evaluated a key prediction on the peripheral-polygenic component – a positive correlation between GWP marker loadings and gene expression in relevant tissues. GWP loadings are indeed correlated with gene expression in relevant tissues. However, correlations are often higher with non-relevant tissues from earlier growth stages or tissues, which is not predicted by within-tissue *trans* regulation for the peripheral-polygenic component (the “omnigenic in space” model). Therefore, these findings suggest that genes expressed in early growth stages, with indirect effects on later traits, are contributing to polygenic variation (“omnigenic in time”). Together, our findings suggest that an extended OM, with regulatory effects both in space and in time, could explain the ubiquitous geometric genetic architecture of complex traits.

## INTRODUCTION

Understanding genetic architecture – the structure of the genotype-phenotype relationship – is a fundamental goal in biology^1–3^. An understanding of genetic architecture is also key to addressing human disease, guiding conservation, and improving livestock and crop species. In plant sciences, genetic architecture is a unifying theme that connects the theoretical understanding of plant evolution and development to applications in plant conservation and breeding^4^. The genetic architecture of yield in crop plants is notably complex because almost any gene affecting growth, development, or environmental response can influence the ultimate yield^5,6^. Genome-wide association studies (GWAS) and linkage mapping have been performed in many plant species to elucidate the genetic architecture of complex traits^5,7^. However, the genetic architecture of yield formation dynamics is much less understood, due to the practical challenge of time-course phenotyping, as well as the conceptual challenges of linking dynamic endophenotypes to each other or end-point phenotypes^8^.

An important outstanding question in genetics is how to integrate statistical quantitative genetics, which provides quantitative predictions of phenotype without mechanistic detail, with molecular and cellular genetics, which provides rich mechanistic detail without generally providing predictions on quantitative variation^9^. The omnigenic model, a new quantitative genomics framework that integrates classical quantitative genetics with molecular and developmental genetics, may contribute to such a unification^10,11^. As in classical models of complex traits, such as Fisher’s geometric model^12^, the omnigenic model posits a geometric genetic architecture with an oligogenic and a polygenic component of allele effects (**Fig. 1a**). Critically, however, the omnigenic model provides a mechanistic rationale for this architecture based on molecular and cellular biology: under the omnigenic model, “core” genes directly regulate the trait and underlie the oligogenic component (i.e. major loci); while “peripheral” genes, which are expressed in the same tissues, but indirectly affect the trait via *trans*-regulation of core genes, underlie the polygenic component. The omnigenic model has spurred vigorous debate in the human genetics literature ^11,13,14^ but in plant systems, which have advantages for tractability, investigations are just beginning ^15,16^.

**Fig. 1:**
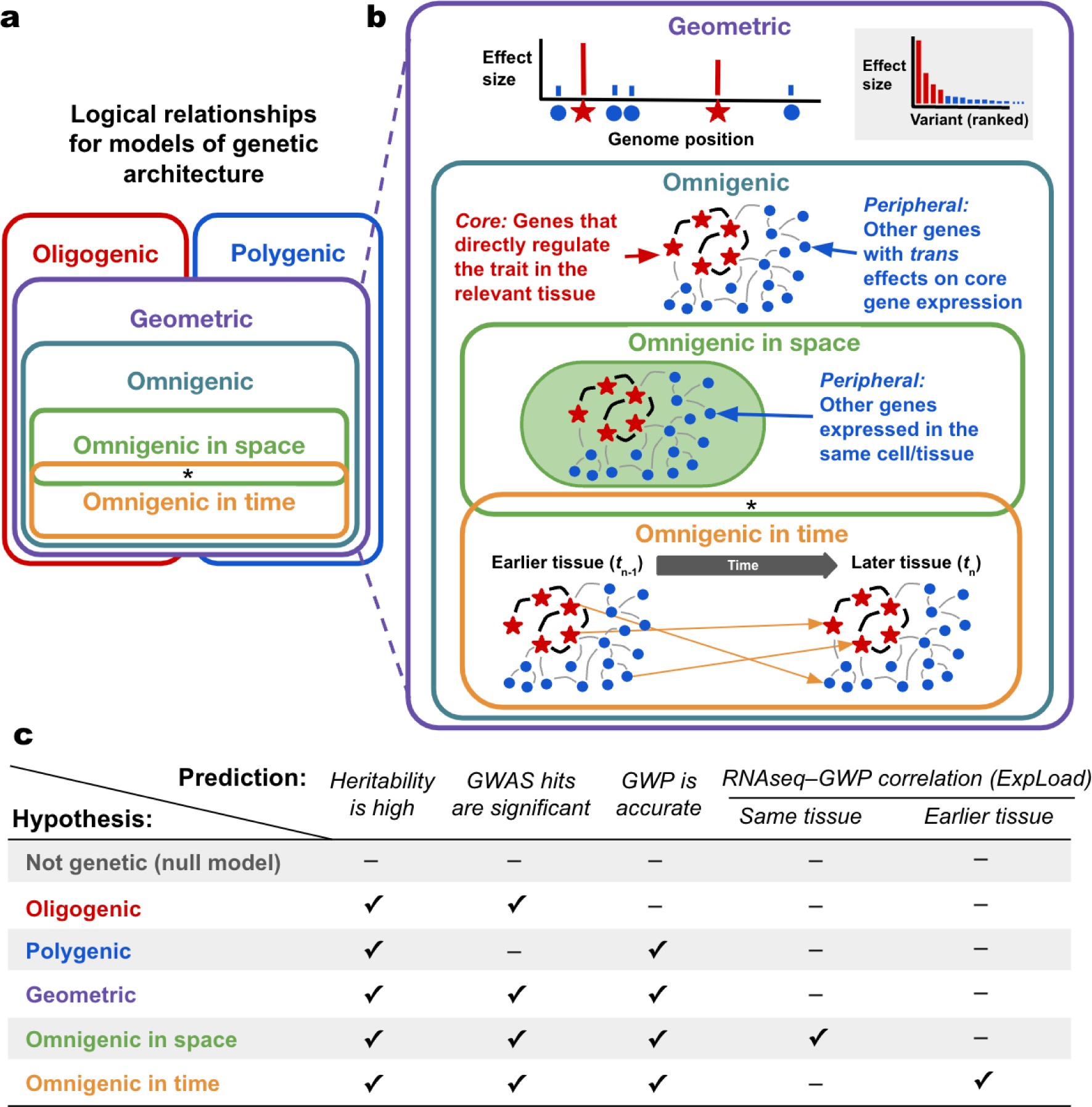
Omnigenic models and framework for testing hypotheses on genetic architecture. **(a)** An Euler diagram summarizing the logical relationship of omnigenic models of the genetic architecture of complex traits versus standard models. (**b**) A detailed description of each model. The *oligogenic* model considers few variants of major effects, while the *polygenic* model considers many variants of small effects. The *geometric* model includes both an oligogenic and polygenic component, consistent with standard theoretical frameworks such as Fisher’s Geometric Model ^12^. Omnigenic models represent a subset of geometric models that posit that variants in core genes (directly regulating the trait) underlie the oligogenic component and peripheral genes (not directly regulating the trait but expressed in the relevant tissue) underlying the polygenic component. We describe the original omnigenic model of Boyle et al. (2017) as *omnigenic-in-space*, as it posits that the peripheral gene network comprises genes expressed in the same cell or tissue (i.e. space) as the core genes. By contrast, we introduce here an *omnigenic-in-time* model, where expression in earlier tissues affects traits in later tissues (which is not mutually exclusive with the omnigenic-in-space model, noted by the asterisk). Red stars: core genes, blue circles: peripheral genes, black lines: regulation among core genes, and gray lines: *trans* regulation by peripheral genes. **(c)** Genetic architecture hypotheses and corresponding predictions for each analysis. The check mark (✓) indicates that the given observation is expected under the given hypothesis, while the hyphen (−) means the given observation is *not* expected. ExpLoad represents the overall correlation between gene expression and marker loading from GWP averaged over each gene.

Plant phenomics promises to elucidate the genetic architecture and molecular basis of developmental and growth dynamics underlying complex traits like yield^17^. Initially, advances occurred in functional genomics (e.g. transcriptomics and metabolomics), which enables the characterization of molecular endophenotypes at unprecedented speed and scale^18,19^. More recently, phenomics of morphological and physiological traits has advanced, particularly with imaging-based high-throughput phenotyping (HTP) platforms suitable for single plant or plot scales^20,21^. HTP from unoccupied aerial systems (UAS) is especially useful for plant biology as it facilitates the characterization of growth and developmental dynamics under ecologically and/or agriculturally relevant field settings and spatial scales.

Biomass sorghum (*Sorghum bicolor* [L.] Moench) provides an excellent system for studying the genetics of yield formation. Unlike in grain crops, where the dynamics of yield formation are observed via destructive measurements, the dynamics of yield formation throughout development in biomass crops can readily be measured via UAS-HTP^21,22^. Sorghum is a short-day plant, which originated in tropical Africa and diversified across tropical and temperate regions across several continents^23^. Photoperiod-sensitive (PS) germplasm provides a basis for high-biomass sorghum since in temperate latitudes PS genotypes delay flowering indefinitely and accumulate massive quantities of vegetative biomass (>20 Mg ha^-1^)^24,25^. The abundant genetic diversity of PS germplasm has been captured in a global biomass association panel (BAP) genomic resource^26^ that has been characterized by whole-genome resequencing^27^.

In this study, we sought to test the omnigenic model, using biomass yield formation in a global sorghum diversity panel. We considered multiple working hypotheses (**Fig. 1a-b**) and developed predictions based on field phenomics, transcriptomics, GWAS, and the genome-wide predictions that could be used to exclude each of these hypotheses (**Fig. 1c**)^28^. The genome-phenome analyses reveal a geometric genetic architecture of biomass yield formation, with distinct oligogenic-core and polygenic-peripheral components, consistent with the omnigenic model. However, the findings suggest that the polygenic-peripheral component is driven, in part, by gene expression in earlier growth stages in tissues not directly related to the trait of interest. This extended omnigenic model goes beyond the original omnigenic model, which proposed that peripheral effects are only due to non-trait-related genes expressed in the same tissue or cell type of the focal trait. Together, our findings suggest that the geometric architecture of complex traits may be explained by the omnigenic model, but that peripheral effects also occur through a variety of processes in tissues that are distant in both time and space.

## RESULTS

### UAS-HTP characterizes genetic variation of traits related to biomass yield formation

To characterize the dynamics of biomass yield formation in global sorghum diversity, we evaluated the panel in replications over two growing seasons and employed UAS-HTP multispectral imaging twice weekly, weather permitting, throughout the growth cycle (**Fig. 2a**). High correlation (*r* = 0.96, *p* < 0.01) between UAS-HTP and manual measured height established the accuracy of the UAS-HTP (**Fig. 2b**). Reduced correlations (*r* = 0.91, *p* < 0.01; Supplementary Fig. 1) later in the season likely reflect characteristics such as lodging or shading by taller genotypes in neighboring plots that are expected for unimproved PS germplasm in temperate field conditions (**Supplementary Fig. 2**). Broad sense heritability (*H*^2^) across blocks (repeatability) was high (0.5–0.9) through most of the growing season (**Fig. 2c**). To complement UAS-HTP, we manually assessed 13 quantitative traits and one qualitative trait, midrib color (which is closely related with lignin content and cellulosic ethanol yield), as a positive control for core gene identification (**Supplementary Table 1 and Supplementary Fig. 3**)^29^. We observed significant genetic variation (*p* < 10^−15^) for all quantitative traits, with the heritable genetic component of variation (*H*^2^) across years ranging from moderate (leaf width: 0.24) to high (flowering time: 0.94) (**Supplementary Table 1**).

**Fig. 2:**
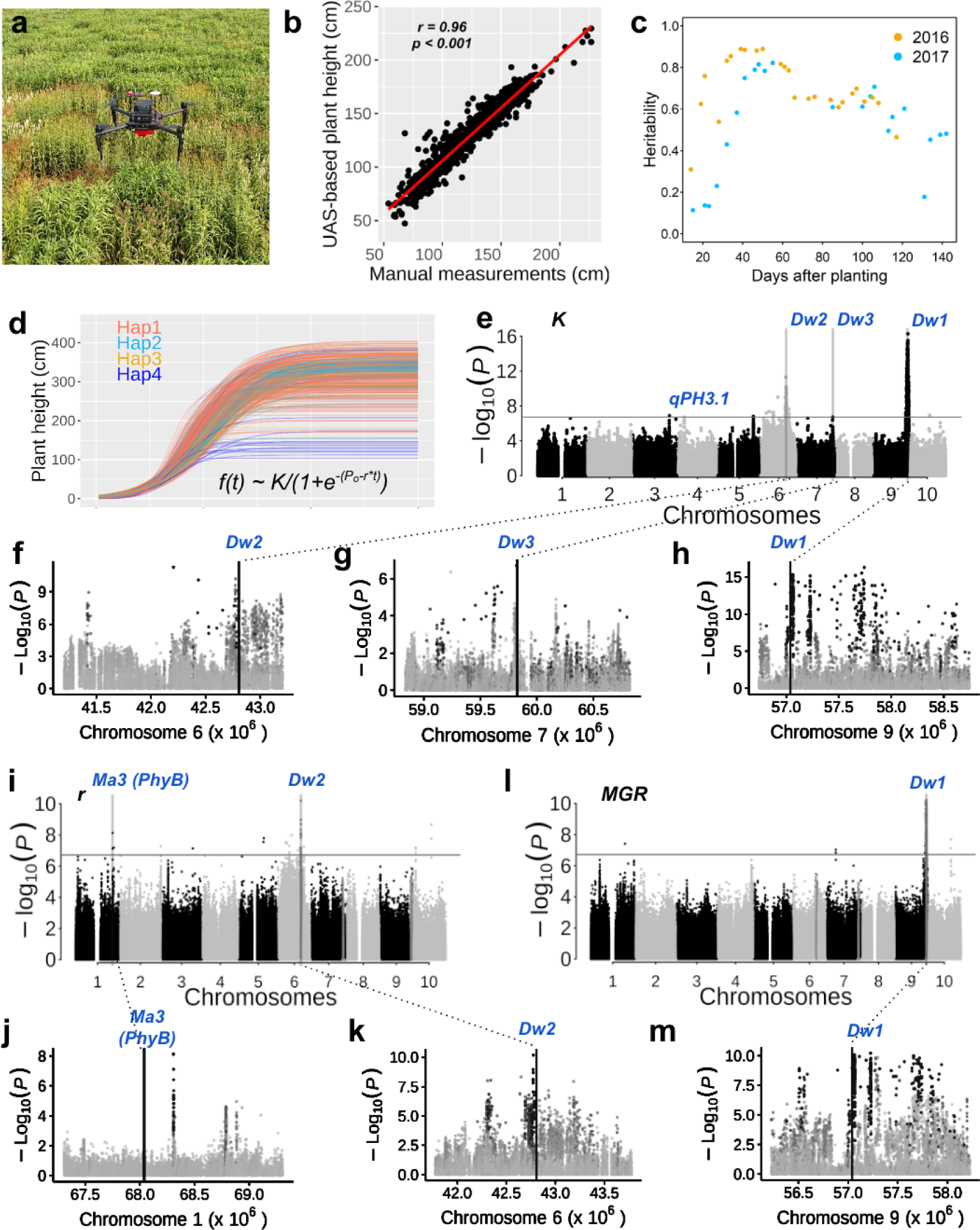
Genome-phenome analyses accurately dissect dynamic genetics of biomass yield formation in global sorghum diversity. (**a**) The UAS-HTP platform collects biweekly field imagery. (**b**) Correlation between UAS-based plant height and manual measurement. Data was collected 41 days after planting in 2016. (**c**) The heritabilities of UAS-based plant height along with sorghum growth. The different colors indicated heritabilities in two years. (**d**) The modeled growth curve for each accession. Each smooth curve means each individual with the equation noted. The colors indicated four haplotypes of *Dw1*, the primary plant height regulator. (**e**) GWAS of growth model parameter *K*, representing the maximum plant height, using a mixed linear model. The known dwarf genes, including *Dw1*, *Dw2*, and *Dw3*, were annotated as red texts. (**f**), (**g**), and (**h**) The zoomed regional association for *Dw2*, *Dw3*, and *Dw1* and linkage disequilibrium between SNPs and the corresponding leading SNPs. (**i**) GWAS of growth model parameter *r*, the instantaneous growth rate. (**j**) and (**k**) Zoomed regional association and linkage disequilibrium between SNPs and leading SNPs for *Ma3* and *Dw2*. (**l**) GWAS of the max growth rate using a mixed linear model. *Dw1* was marked as blue text. (**m**) The zoomed regional association and linkage disequilibrium between SNPs and leading SNPs for *Dw1*.The linkage disequilibrium (*r*^2^) between each SNP within the region and the leading SNP is denoted with the grayscale.

### Modeled growth dynamics underlying biomass yield formation

We used the UAS-HTP measurements and the final manual plant height measurements to model the growth dynamics of plant height for each individual, using a simple non-linear logistic model with three parameters: *K*, the maximum plant height; *P*_o_, the time of onset of exponential growth; and *r*, the instantaneous proportional growth rate (**Supplementary Fig. 4**). We used the averaged model parameters (*K*, *P*_o_, and *r*) using data from two years to generate plant height growth curves, which showed that haplotype 4 of *Dwarf1* (*Dw1*) had the shortest plant height and lowest growth rate (**Fig. 2d, Supplementary Fig. 4-5**). The modeled height for each genotype, *K*, ranged substantially (79–414 cm; Fig. 2d) and substantial variation existed among accessions for the daily growth rates (cm/day) across the growing season (**Supplementary Fig. 4c**). From the growth rate model curve, we extracted four parameters that characterize the differences in plant growth curves: maximum growth rate (*MGR:* 2.8 - 12.9 cm/day), date of *MGR* (*d_MGR_*: 28 - 59 days after planting), average growth rate (*R_a_*:1.6 to 4.2 cm/day), and duration of the high growth rate ranges (*L-days*: 13 to 54 days) (**Supplementary Fig. 4c and Supplementary Table 2**).

Significant (*p* < 10^−16^) genotype differences existed for all parameters, and we observed a significant effect (*p* < 10^−3^) of the environment on phenotypic variation for all parameters other than *MGR* (**Supplementary Table 2**). All modeled growth parameters were highly heritable, with broad-sense heritabilities ranging from 0.54 (*R_a_*) to 0.93 (*MGR*). We evaluated potential yield components with correlation analysis and found that many were highly correlated in the expected direction (*r* = −0.97–0.99) (**Supplementary Fig. 6a-f and Supplementary Data File 1**). Biomass yield increased with plant height, but the relationship was nonlinear, plateauing after ∼350 cm (**Supplementary Fig. 6b**). Fresh biomass was more highly correlated (*r* = 0.64, *p* < 0.001; **Supplementary Fig. 6c**) with leaf length than plant height (*r* = 0.64, *p* < 0.001; **Supplementary Data File 1**). Flowering time is the major event during which plants shift from vegetative to reproductive growth, and *MGR* occurs an average of 14 days before flowering time (**Supplementary Fig. 6d**).

### GWAS of manually-collected phenotypes validates the genome-phenome resource

We first investigate the putative oligogenic-core genetic components of biomass yield formation manually collected phenotypes, conducting GWAS using 5,215,082 single nucleotide polymorphisms (SNPs) with a mixed linear model (MLM). A total of 107 genome-phenome associations were identified, with evidence of several putative pleiotropic loci (**Supplementary Data File 2 and Supplementary Fig. 7-14**). A well-characterized gene, the *Dry* locus, was identified for midrib color (a proxy for stem and leaf juiciness) with the strongest associated SNP (hereafter called “leading SNPs”) on chromosome 6 (S06_50893225; *p* < 10^−7^) localized 2.4 kb upstream of the cloned *Dry* transcription factor Sobic.006G147400 ^29^ (**Supplementary Fig. 3b**).

A notable novel association (*qPH3.1*; MLM *p* < 10^−7^) for plant height on chromosome 3, which was improved by including *Dw1* and *Dw2* as covariates in MLM or using a multi-locus mixed model (MLMM) (**Supplementary Fig. 10, Supplementary Fig. 11a, and Supplementary Fig. 11b**). Two leading SNPs (S03_61595713, S03_61860987) were under strong linkage disequilibrium (LD) (*r*^2^ = 0.88; *p* < 0.001), suggesting that they tagged the same underlying causal variant of *qPH3.1*. Searching through the associated region, we found that SNP S03_61595713 is localized in a putative AP2/B3-like transcriptional factor family protein gene (Sobic.003G281100; *SbVRN1-B3*) (**Supplementary Fig. 11c**). Its ortholog in *Arabidopsis, VRN1* (*AT3G18990*; 42% similarity), represses *FLC* to activate the floral transition^30^. Eight haplotypes were identified for *SbVRN1-B3* in our population and accessions with haplotype 7 were significantly shorter (Tukey HSD, *p* < 0.01) than those with other haplotypes (**Supplementary Fig. 13**). Taken together, these results suggest *SbVRN1-B3* is a strong candidate for the dynamic growth QTL *qPH3.1*.

### GWAS and GWP decompose dynamic core-oligogenic components of biomass yield formation and peripheral-polygenic effects on biomass traits

To elucidate the dynamic effects of the putative oligogenic/core genes, we conducted MLMM GWAS for the modeled growth parameters (**Fig. 2e-2m**). As expected, we observed strong precise associations near known dwarfing genes *Dw1*, *Dw2*, and *Dw3* for the growth curve parameter *K* (**Fig. 2e-h**). Overall, we identified five associations for *P*_o_ and 11 associations for *r* (**Fig. 2i and Supplementary Fig. 14**). Two of the 11 associations for *r* were colocalized with known genes, *Ma3* (*PhyB)* and *Dw2* (**Fig. 2i-k**). *Dw2* was associated with three growth parameters: *K*, *P_o_*, and *r*. The association (S01_68306822) on chromosome 1 for both *r* and *P*_o_ was localized upstream (∼27 kb) of *Ma3* (*PhyB)* (**Fig. 2j**)^31^. *Dw1* was identified for three parameters, including *R_a_*, *d_MGR_*, and *MGR* (**Fig. 2l, and Supplementary Fig. 14**), and *Dw2* was identified for average growth rate and *d_MGR_* and *Ra* (**Supplementary Fig. 14)**. One common SNP (S01_63035838) on chromosome 1 was identified as a marginal association for *R_a_* (*p* < 10^−6^) and *MGR* (*p* < 10^−7^) (**Supplementary Fig. 14**).

To identify the oligogenic-core components of plant height dynamics, we performed GWAS for plant height growth curves and growth rate curves using the MLMM, which can identify the three major QTLs/genes for plant height (**Fig. 3a and b**). We identified three persistent QTLs (*Dw1*, *Dw2*, and *qPH3.1*) and two transient QTLs (*DqPH1.1* and *DqPH2.1*) for plant height growth curve (**Fig. 3a**), while nine dynamic QTLs, including *Dw1*, *qPH3.1*, and *Dw2*, were associated with the growth rate curve (**Fig. 3b**). *Dw1* was identified as the earliest, followed by *qPH3.1*, then *Dw2* (**Fig. 3a**). Additionally, two transient QTL (*DqPH1.1* and *DqPH2.1*) were mainly identified around 60 DAP (**Fig. 3a**). Only transient QTL for dynamic growth rate were identified, but earlier than transient QTL of the growth curve (**Fig. 3b**), reflecting high variance of growth rate traits around 50 DAP, and a delay (∼10 days) of the result of the growth rate change. Transient QTL *DqGR9.1* (S09_59245434) was downstream (85 kb) of putative flowering time gene *SbELF3* (Sobic.009G257300) ^32^ (**Table S4**).

**Fig. 3:**
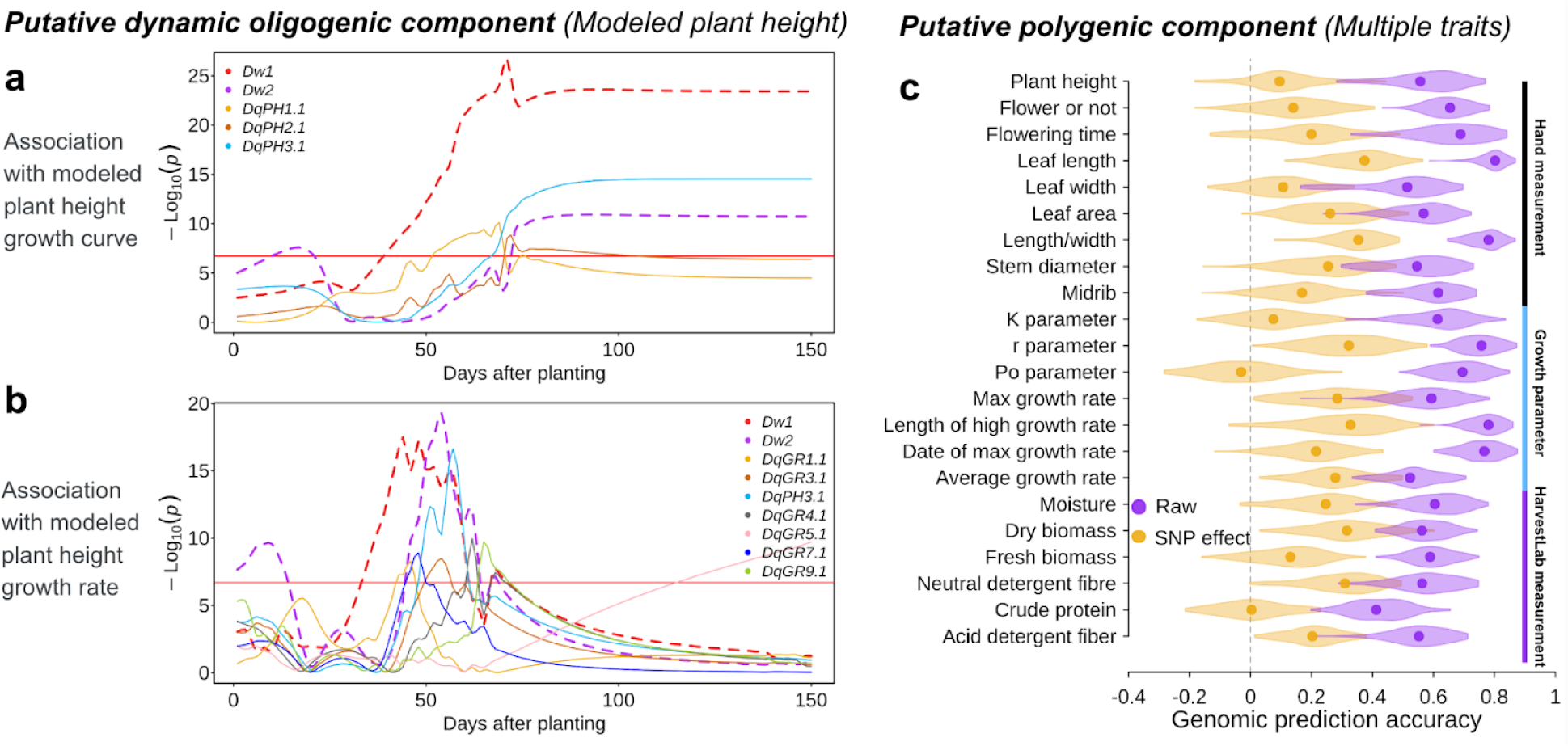
Decomposing the dynamic oligogenic-core and polygenic-peripheral components. The trajectories of multi-locus mixed model association significance of (**a**) modeled plant height (growth curve) and (**b**) modeled plant height growth rate at daily step (Fig. 2d-e). Different colors indicate different QTLs (*Dq*). The dashed lines represent associations that colocalize with known genes (*Dw1* and *Dw2*), while the thin lines represent other associations. Novel dynamic trait associations are named as *Dq*+*trait*+*order on the chromosome*, the number means the chromosome and the order on the chromosome. **(c)** Genomic predictions that account for the oligogenic-core component of variation reveal the putative polygenic-peripheral component. Genome prediction accuracy for 22 biomass traits or growth model parameters with violin plot color indicating contrasting GWP models: “Raw” (purple) indicates genomic prediction was conducted using raw phenotype (BLUP score); “SNP effect” (yellow) indicated training the genomic prediction model using treated phenotype by removing the effect of associated SNPs (represented by the lead SNP), so variation accounted for by oligogenic-core genes would be excluded. The genome prediction accuracy was estimated using 5-fold validation with 100 permutations.

Under geometric models of genetic architecture, including omnigenic models (**Fig. 1a-b**), the oligogenic component is expected to be captured by GWAS while the polygenic component is to be captured by a GWP once the contribution of oligogenic variation has been removed (**Fig. 1c**). Therefore, to further decompose the oligogenic and polygenic genetic components of phenotypic variation for biomass traits, we compared the accuracy of GWPs using either a standard GWP model (“Raw” model) or including the putative oligogenic component from GWAS as covariates in the GWP model to isolate the polygenic component (“marker effect”) (**Fig. 3c**). With the raw model, mean prediction accuracies were high, with a range from 0.40 (crude protein content) to 0.80 (leaf length) (**Fig. 3c**), presumably reflecting that both the oligogenic and polygenic components are contributing to prediction accuracy. By contrast, After accounting for the putative core-oligogenic component (“Covariates” model), GWP accuracies were substantially reduced compared to the raw model (−0.08 for leaf width to 0.53 for NDF), but greater than zero for most traits except leaf width (**Fig. 3c**), which would correspond to the peripheral-polygenic component of the omnigenic model.

### Correlations of tissue-based gene expression from RNAseq and gene loadings from GWP (ExpLoad) carry signatures of omnigenic effects

The omnigenic model states that most genes expressed in trait-relevant tissues contribute to the genetic variation of complex traits via the peripheral-polygenic component^10^. Under that proposition, we reasoned that gene expression in trait-relevant tissues based on RNA-seq should be positively correlated with genome-wide marker loadings from the GWP (**Fig. 1, Supplementary Fig. 15**). Marker loading varied across genomes and major associated loci from GWAS showed high marker loading (**Supplementary Fig. 15**). (Note, we use the term “marker loading” rather than the more common term “marker effects’’ to clarify that SNP used as markers tag causative variation but are not necessarily causative). Since expression data are per-gene, we sought to compare this with a per-gene estimate of the contribution to quantitative trait variation from the GWP. Thus, for each trait and each gene, we averaged marker loadings from the GWP over the given gene, including exons, introns, and promoter region (5 kb upstream) to calculate the “gene loading”. Then we calculated the relationship between gene loading for the biomass traits and RNAseq across 844 RNAseq data sets in 19 tissues^33^ (**Fig. 1b**). We refer to this metric which captures the relationship between observed tissue-based gene expression and estimated polygenic effects per gene (*R*_Gene loading, gene expression_) as “expression loadings” (ExpLoad).

Overall, the ExpLoad estimates were positive across all tissue and trait combinations (0.02–0.07; **Fig 4a**), suggesting that higher expressed genes have a higher contribution to polygenic trait variation, consistent with the idea of peripheral effects under an omnigenic model. Interestingly, some, and the highest ExpLoad on traits were not relevant tissue that would have been expected under the omnigenic-in-space model (**Fig. 4a, Supplementary Fig. 16a**). Overall, reproductive/late tissues had low ExpLoad, and vegetative/early tissues had high ExpLoad, regardless of the trait (**Fig. 4a-c**). To better characterize the phenomena, we looked in more detail at two traits, leaf length, and plant height, where an obvious relevant tissue for gene expression could be defined (“leaf” and “shoot”, respectively; **Supplementary Fig. 17**). For leaf length, leaf expression had the second highest ExpLoad (*ρ* = 0.055) of 19 tissue types (**Fig. 4b**). For plant height, the *a priori* relevant tissue, shoot, had a moderate ExpLoad (*ρ* = 0.044), ranked 8th across 19 tissues, while leaf again had the second highest ExpLoad (*ρ* = 0.056) (**Fig. 4c**). Notably, for both traits, the highest ExpLoad was observed in roots (*ρ* = 0.059) (**Fig. 4b-c**). While the positive ExpLoad across many tissue-trait combinations is consistent with the existence of omnigenic effects, the high ExpLoad for tissues that would be considered non-relevant under the omnigenic-in-space model suggests that this model is not sufficient to explain the genome-phenome relationships.

**Fig. 4:**
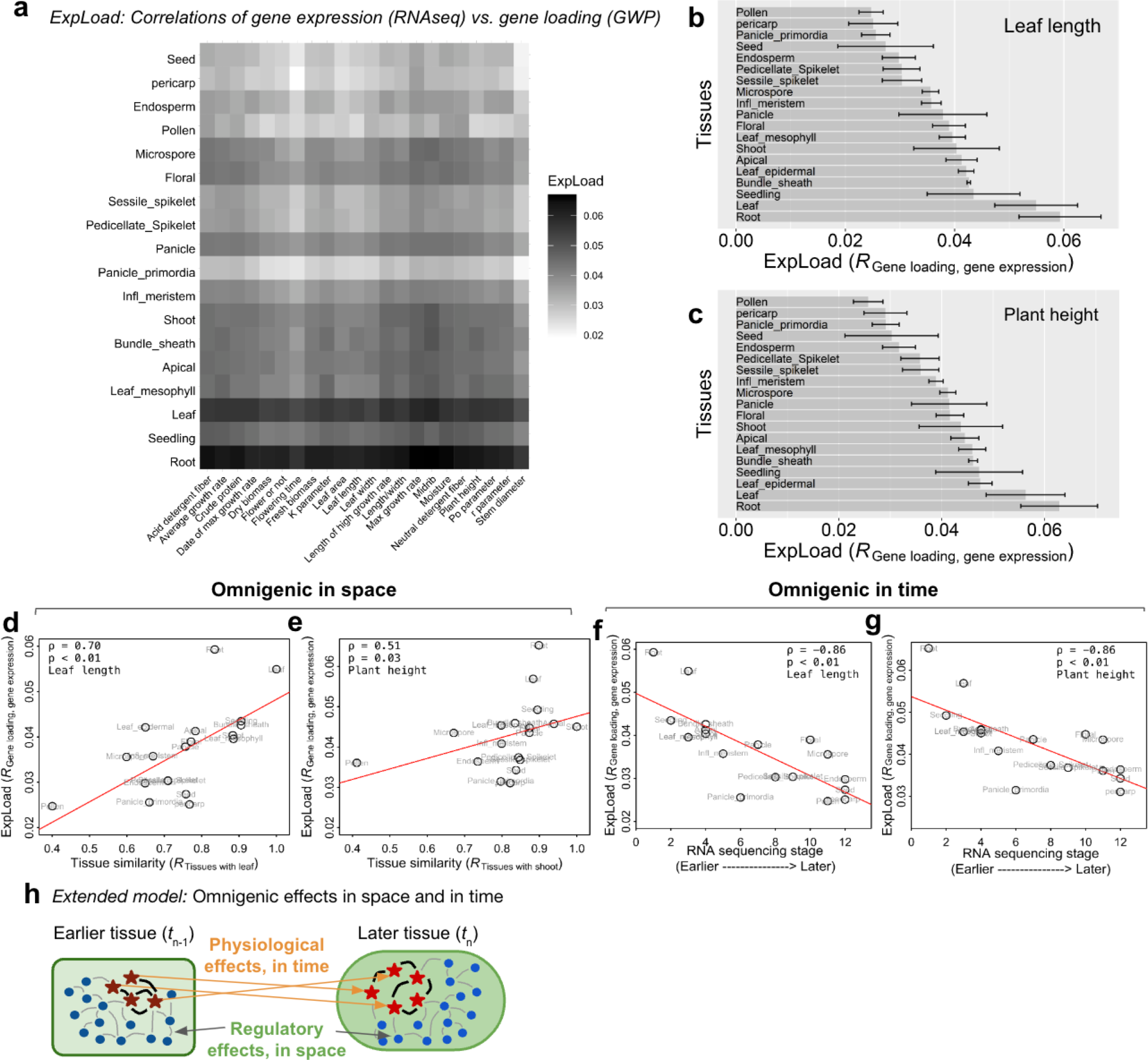
Testing the omnigenic model based on predicted correlations of polygenic effects and tissue-level gene expression. (**a**) The correlation coefficient heatmap of RNAseq from different tissues and gene loading from genome prediction for different traits. The “gene loading” was calculated using the marker loading from the genome prediction for SNPs located in a given gene and upstream 5 kb (regulatory region) was used. The order of the tissues from bottom to top represented the order the tissue arises. (**b**) The sorted correlation coefficient for gene loading of leaf length and RNAseq crossing different tissues. (**c**) The correlation coefficient for gene loading of plant height and RNAseq crossing different tissues. The error bars (+/− standard deviation) were calculated using RNAseq replicates from the same tissue. (**d**) and (**e**) the correlation of correlation ExpLoad and tissue space for leaf length and plant height. Tissue space was calculated as the correlation coefficient of RNAseq between other tissues and relevant tissue of the trait. (**f**) and (**g**) the correlation of correlation between ExpLoad and times for leaf length and plant height. Tissue time is defined by the order of tissue samples in the sorghum life cycle. (**h**) A model integrated with omnigenic in space and time. Black-red and red stars mean core genes in earlier tissue and later tissues. Dark blue and blue stars mean peripheral genes in earlier tissue and later tissues.

### Relationship between ExpLoad and tissue similarity or tissue stage distinguish omnigenic effects in space versus time

To more specifically distinguish omnigenic-in-space^10^ and omnigenic-in-time (**Fig. 1b**) effects we sought quantitative metrics of tissue similarity and growth stage, respectively, which could be correlated with ExpLoad (**Fig. 1c**). We address each of these in turn:

First, we reasoned that under the omnigenic-in-space hypothesis, (1) similarity within “tissue space” could be estimated by the overall correlation between gene expression in the *a priori* trait-relevant tissue with other tissues and (2) this tissue similarity metric should itself be positively correlated to ExpLoad for a given tissue. This is because the omnigenic-in-space model posits that trait variation is affected by the expression of non-trait-relevant (peripheral) genes in the same tissue as the trait-relevant (core) genes that directly regulate the traits. Thus, to test the omnigenic-in-space hypothesis for leaf length we estimated the correlation, *ρ*, of tissue similarity with leaf (*R*_Tissue with leaf_) with ExpLoad for each tissue (**Fig. 4d**); and to test the hypothesis for plant height we estimated the correlation, *ρ*, of tissue similarity to shoot (*R*_Tissue with shoot_) (**Fig. 4e**) with ExpLoad for each tissue. For both leaf length (*ρ* = 0.70, *p* < 0.01; **Fig. 4d**) and plant height (*ρ* = 0.51, *p* = 0.03, **Fig. 4e**), a significant correlation was observed, consistent with the predictions under the omnigenic-in-space hypothesis for these traits. Further, most traits showed a significant association between tissue similarity and ExpLoad (**Supplementary Fig. 17**).

Secondly, we reasoned that under the omnigenic-in-time hypothesis, staged tissue should be negatively correlated to ExpLoad for a given tissue. This is because an omnigenic-in-time model would posit that early-expressed genes affect later traits, including those in other tissues. Overall, we observed highly negative correlations between staged tissue and ExpLoad (**Fig. 4f and 4g; Supplementary Fig. 18**). For leaf length, the time of RNAseq was highly correlated (*ρ* = −0.86, *p* < 10^−5^) with ExpLoad (**Fig. 4f**). Similarly, for plant height, there was a high correlation between staged tissue and ExpLoad (*ρ* = −0.86, *p* < 10^−5^) (**Fig. 4g**). Thus, observations on the genome-phenome relationships are consistent with the hypothesis that both omnigenic-in-space and omnigenic-in-time effects are acting on trait variation for complex traits in global sorghum diversity (**Fig. 4h**).

## DISCUSSION

The formation of yield is a dynamic physiological process resulting from genetic and environmental interaction, underpinned by complex gene regulatory networks that change overgrowth and development^34^. In this study, we used high-throughput genome-to-phenome analyses of diverse global sorghum – integrating whole genome resequencing, transcriptome analysis, and field phenomics – to elucidate the genetics of dynamic biomass yield formation in the field. The high accuracy and heritability of UAS-HTP measurements (Fig. 2b-c) established this as a reliable approach for large-scale genetic variation of dynamic growth of sorghum^21,35–37^. Further, The precise identification of cloned natural variants (Fig. 2e-m) validated the genome-phenome data set and modeling. Overall, the genome-phenome relationships (Fig. 2-4) support the original omnigenic model, which focused on *trans* regulation of core genes and peripheral genes to impact the phenotype^10,11^, while also expanding to new dimensions beyond this original model (**Fig. 1**).

The original omnigenic model was proposed to explain observations of geometric genetic architecture and missing heritability for human disease^10^. Since this model postulates that tissue-specific gene expression is the critical determinant of trait expression in diseased cell types, previous studies assumed that genes do not contribute to heritability unless they are expressed in the trait-relevant tissues^11^, and genes in the relevant pathway (core genes) or expressed in relevant tissue (peripheral genes) contribute most to heritability^10^. Two key observations of the human studies that support the omnigenic model were: (1) those large-effect genetic variants identified in GWAS are central pathways of the relevant tissues, and (2) peripheral gene expression in the relevant tissue plays the most important role in determining the polygenic variation of a disease trait in human^10^. In our study, GWAS signals (**Fig. 2e-m**) captured large effect variants that could be traced back to central pathways for the given trait, consistent with observation (1) in humans^38,39^. Consistent with observation (2) from human studies, GWAS-identified variants only captured part of the heritability (**Fig. 2, 3a-b**), while whole-genome prediction captured most of the remaining variation (**Fig. 3c**), confirming a geometric genetic architecture for each of the study’s traits.

In our study, the polygenic model was able to account for much of the variation in most traits (**Fig. 3c**). This may be an underestimate because core gene effects on phenotypic variation are estimated first, and part of the peripheral effects may be absorbed due to genome-wide LD^40^. Part of the missing heritability^41^ we observed (**Fig. 2c**) may be “hidden” heritability due to minor effect loci that fall below GWAS threshold^38^, which in our study would represent the polygenic component of yield formation (**Fig. 1 and 3c**). Under the omnigenic model, the hidden heritability would be largely due to variation in peripheral genes expressed in the relevant tissue^10^. However, we observed that gene expression in unrelated tissues had the strongest correlation with the polygenic component, particularly, gene expression at early development stages (**Fig. 4f-g**). This suggests an omnigenic-in-time mechanism (**Fig. 1**) where gene expression at earlier developmental stages combines to be major determinants of phenotypes of later development stages (**Fig. 4h**). For example, root expression of genes is particularly predictive of polygenic effects on other tissues (**Fig. 4d-g**). Given the importance of roots for the uptake of water and nutrients, it seems plausible that many traits in above-ground tissues would be affected by genes expressed early in root development. A previous study showed that artificial selection during domestication reshaped the architecture of the roots, but no colocalized associations were identified for roots and ground-above traits^42^, suggesting a shared peripheral gene network between roots and ground-above traits. Our findings provide genomics evidence of the persistent impact of root gene expression on plant traits.

From a theoretical standpoint, it seems likely that complex traits are affected by earlier gene expression in developmentally and/or physiologically important tissues that are distant from the tissue expressing the trait of interest, as posited in the omnigenic-in-time model. For developmental traits, such as biomass yield formation, many biological processes affect yield formation, and many studies have shown that non-related tissue affects each other in plants^42^. For instance, hormones by definition act in tissues other than those in which they are synthesized^43^ so hormone biosynthesis and transport genes expressed in “non-relevant” tissue would contribute to heritability. Our study suggests that, at least in the case of biomass yield formation, indirect effects of genes expressed earlier in growth and development are equally important contributors to the peripheral gene network (**Fig. 4d-e vs. Fig. 4f-g**).

Early-stage transient expression of large effect genes, which could be considered the oligogenic-core component, may explain part of the hidden heritability for endpoint traits, because their effects may be dampened over time. For instance, some transient height QTL (**Fig. 3a-b**) had reduced effect sizes over time and contributed little to final height (**Fig. 2e**), presumably due to later-expressed height QTL swamping out their effect. Interestingly, these findings are consistent with Sinott’s 1937 hypothesis “that genes control[ing] the rates of growth processes” across “successive increments of growth” leads to the “geometrically progressive character of genic effects” (i.e. geometric genetic architecture)^44^.

The effect of early-expressed genes on later fitness traits is central to theoretical debates on evolutionary development models, such as the contribution of“developmental burden” underlying von Baer’s Third Law ^45^. Similarly, the idea that many or most early expressed genes will affect final yield is widely held and longstanding in the plant breeding community. For instance, Allard’s 1960 *Principles of Plant Breeding* states that “mutations, particularly those … expressed early in the life of the individual, can produce a syndrome of effects“^46^. While this concept is not currently formalized in genomic selection methods, new prediction frameworks that incorporate phenomic data (e.g. transcriptome, metabolome) would make it possible to do so^47^, particularly to generate and select polygenic variation in elite breeding germplasm where major effect variants in core genes are already fixed^48^. Thus, new genomic selection approaches based on the omnigenic model could contribute to improved food security and climate resilience. To further test the omnigenic-in-space and omnigenic-in-time models, and estimate their relative contributions, cell-type specific ExpLoad predictions for each model could be tested with single-cell RNA-Seq in relevant and non-relevant tissues^49^.

In this study, we have bridged the omnigenic model from human genetics to plant systems, by formally decomposing the oligogenic-core and peripheral-polygenic components of variation. We validate several key predictions of the model, which are consistent with numerous observations of geometric genetic architecture from decades of genetic studies in plants, but connect this with gene expression of the polygenetic component. At the same time, we have key observations that extend the omnigenic model, particularly including the importance of gene expression at early developmental stages. Under this extended omnigenic-in-time model, a substantial portion of the polygenic component of endpoint traits (e.g. yield, disease severity) would be “reverberations” of the transient oligogenic component (i.e. early expressed QTL; e.g. **Fig. 3a**) from related, and even unrelated traits and tissues, earlier in growth and development. Overall, our genome-phenome analysis supports the hypothesis that geometric genetic architecture emerges as a consequence of transient gene expression over growth and development.

## MATERIALS AND METHODS

### Plant materials and field design

Three hundred sixty-three accessions from the Bioenergy Association panel^26^ were for genome-phenome analyses. The germplasm was evaluated at the Kansas State University Ashland Bottoms Research Farm (39°08’25.5“N, 96°37’54.5“W) and planted on June/17th, 2016, and May/30th, 2017. The experiment was designed with the randomized complete block design in a four-row plot with two replicates each year. Two commercial hybrid cultivars, Pacesetter (Richardson Seeds Inc., TX) and Blade ES5200 (Ceres Inc., CA), were used as the border between and around two replicate blocks.

### Field phenomics image collection and image processing

Field imagery was collected using the MicaSense RedEdge multispectral camera (MicaSense Inc., United States) carried on a quadcopter UAS (DJI Matrice 100, DJI, Shenzhen, China). Two times collections per week were performed, as weather allowed. The evaluation of the UAS flown was at 21 meters, and the 70% overlap between the continued two images was to promise the mosaic of the image. The image was processed using our published method^21^. The data quality was checked by calculating the heritability and the correlation between UAS data and manual measurements. The heritability was calculated as the repeatability by using the UAS data from two blocks with the heritability package^50^. The correlation was performed using the cor.test function in R. The underlying imagery used in this study has been integrated into a published data set^51^.

### Manual phenotypic measurements

Plant height at the different stages of the growing season was measured in centimeters (cm) using a barcoded wooden stick and scanner. The width of the third stem node from the root was measured using digital calipers in millimeters (mm). Leaf length (mm) and leaf width (mm) were measured using the barcoded ruler and the leaf length and leaf width were recorded using a digital scanner. Flowering time was collected as the date when 50% plant flowers in the plot and converted to days after planting. Since some germplasms are sensitive to the photoperiod, they can not flower. We converted the feature as a category trait: flowering or not flowering. The color of the midrib was recorded as white and brown (Supplementary Fig. 3a). Fresh biomass per plot was measured using the central two rows in each plot using the John Deere 5460 forage harvester with a custom load cell and converted to kg/m^2^. The chemistry content, including crude protein, acid detergent fiber, neutral detergent fiber, and moisture, was measured using HarvestLab (John Deere, Illinois), and the dry biomass was calculated using the fresh biomass and moisture content.

### Phenotypic data analysis

The best linear unbiased prediction (BLUP) value for each quantitative trait was calculated using a mixed linear model with the lmer function in the lme4 package ^52^

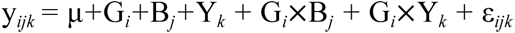

y*_ijk_* is the phenotypic data of *i*^th^ genotype under *j*^th^ block and *k*^th^ year, μ is the overall mean, G*_i_* is the random genotype effect, B*_j_*is the random block effect and Y*_k_* is the random environmental (year) effect, ε*_ijb_* is the residual. The heritability was calculated using the variance from the model

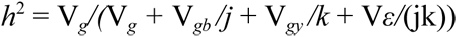

V*_g_* is the variance of genotype V*_gb_* is the variance of interaction for genotype and block, V*_gy_* is the variance of interaction for genotype and year, V*_ε_* is the variance of residual. To test the effect of block and environment effect (year) on phenotype, general linear regression was run using the *lm* function in R with the following model:

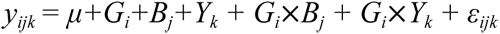

### Plant height growth modeling

Plant height growth of growth seasons was modeled with the equation *f(t)* = *K*/(1*+e^-(Po+r*t)^*) using the *nls* function in R. In the initial model, the *P*_o_ was set as the first measurement from UAS-data, *r* was set as −0.21, and the *K* was set as the plant height (hand measurement) at harvesting. The *t* is days after planting. The equation was modeled using the UAS data for two years data, respectively. The raw data higher than the final plant height (hand measurement) was removed from the dataset, it could be caused by the complex field conditions.

To merge two years of data, the averaged value for three parameters was calculated using the two growth models from two years of data with the lme4 package and was used to fit the growth curve. The absolute growth height per day (*R_pd_*) was extracted using the derivative of the growth model for each genotype. The max growth rate (MGR) and the days after the plant growth rate reached the max growth rate (*d_MGR_*) were extracted from the growth rate curve. The time length of keeping high-speed growth was determined *as L-days* = abs(*dap_MGR1* - *dap_MGR2*). Where *dap_MGR1* is the first time the growth rate reaches *½ MGR* and *dap_MGR2* is the second time the growth rate reaches *½ MGR*. The average growth rate (*R_a_*) for each accession was calculated as 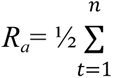 *Rpd*, where n is the days from planting to *d_MGR_*.

All the above growth parameters were extracted from the models for two years, and the BLUP value was calculated using the two-year data.

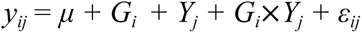

where *y_ij_* is the phenotypic data for the *i^th^* genotype under *j^th^* year, *μ* is the overall mean, *G_i_* is the random genotype, *Y_j_* is the random year. The heritability was calculated using the variance from the model

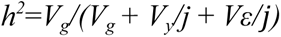

*V_g_* is the variance of genotype, *V_y_* is the variance of between two years, *V_ε_* is the variance of residual. To test the effect of block and year on phenotype, regression was run using the *lm* function in R. The model

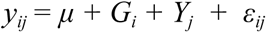

The *ANOVA* function in R was used to extract the effect of each term on phenotype.

### Genome-wide association study and Candidate Identification

Whole-genome sequencing data from the TERRA-REF project was used as genotypic data (https://datadryad.org/stash/dataset/doi:10.5061/dryad.4b8gtht99). SNP data from the whole-genome sequencing data were filtered with minor allele frequency (0.01) and biallelic SNP using VCFtools^53^, and 5,215,082 high-quality SNPs were kept for GWAS in the present study. LMM for GWAS was run with the Genome Association and Prediction Integrated Tool (GAPIT)^54,55^, the model.selection = TRUE, and the kinship was calculated using the default setting. The threshold was set as -log10(1/snp_number). MLMM for GWAS was run with nbchunks = 2 and maxsteps = 10, and results were based on EBIC^56^. LD between SNPs was calculated using the LD function from the genetic package^57^. Gene haplotype was built using the pegas package^58^. Multiple comparisons were conducted using the Tukey test with the *TukeyHSD* function from the agricolae package^59^.

### Genome-wide prediction and dissection of the polygenic component

Genome-wide predictions were conducted using the ridge regression mixed model in the rrBLUP package^60^. Variation in predictive accuracy was estimated using five-fold validation, and the genomic prediction was repeated 100 times by random sampling. To check the impact of identified associations on the prediction accuracy and variation captured by them, the following analyses were performed. First, we removed the major association SNP by around SNP (±50kb) from the leading SNP in the genotypic dataset, then re-run the genomic prediction using the same method as above. Second, we removed the effect of major association and redo the genomic prediction. We removed the effect of major loci by fitting a model:

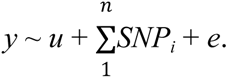

In the model, *u* is the population means, *n* is the number of associations, *SNP_i_* is the *i*th association identified by GWAS, and *e* is the residual. The residual (*e*) from the model was extracted from the model as phenotypic data for the following genomic prediction analysis using the same method as above.

### Omnigenic model test

The gene loading for each trait was averaged over the marker loading within −5 kb of the gene (putative regulatory region) and gene body, and the marker loading was calculated using the rrBLUP using all phenotyped accessions^60^. The gene expression was extracted from a publicly available RNA-seq dataset across 19 tissues (https://zhenbinhu.shinyapps.io/Transcriptome) ^33^. The correlation between gene expression and gene effect was calculated using the *cor.test* function with method = “spearman’’ in R.

## ACKNOWLEDGMENTS

This study was supported by “TERRA-REF: A Reference Phenotyping System For Energy Sorghum” funded by the US Department of Energy’s Advanced Research Projects Agency-Energy (ARPA-E). Data analysis was carried out on the Beocat High-Performance Computing cluster at Kansas State University. The authors declare no competing interests.

## AUTHOR CONTRIBUTIONS

G.M., J.P., and Z.H. designed and conceived the experiments. Z.H. conducted the field experiment design and analyzed the genetic and genomics data. Z.H. and S.M. collected the phenotype data. X.W and J.P. analyzed the imagery for phenotyping. Z.H., J.P., and G.M. wrote the manuscript. All authors reviewed and approved the manuscript.

## Notes

### Competing Interest Statement

The authors have declared no competing interest.

