## Supplemental Tables and Figures for "An extended omnigenic model explains genome-phenome relationships for complex traits in global sorghum diversity"

**Supplementary Table 1 Variation of biomass-related morphological traits**

| **Trait** | **Range** | **Mean** | **Genotype** | **Year** | **Block** | **G**$\times$**Y** | **G**$\times$**B** | ***h^2^*** |
| --- | --- | --- | --- | --- | --- | --- | --- | --- |
| Fresh biomass (kg/m^2^) | 9.7-38.6 | 24.7 | <10^-15^ | <0.01 | <0.01 | <0.01 | <0.01 | 0.75 |
| Dry biomass (kg/m^2^) | 5.1-11.3 | 8.2 | <10^-15^ | Ns | <0.01 | <0.01 | <0.01 | 0.55 |
| Moisture (%) | 46-76 | 65.5 | <10^-15^ | <0.01 | <0.01 | NS | 0.03 | 0.86 |
| Plant height (cm) | 109-442 | 330 | <10^-15^ | <0.01 | <0.01 | NS | NS | 0.92 |
| Flowering time | 50-124 | 83 | <10^-15^ | <0.01 | <0.01 | <0.01 | <0.01 | 0.94 |
| Leaf length (mm) | 521-1278 | 902 | <10^-15^ | <0.01 | NS | <0.01 | <0.01 | 0.91 |
| Leaf width (mm) | 61-105 | 74 | <10^-15^ | <0.01 | <0.01 | <0.01 | 0.03 | 0.24 |
| Ratio leaf length/width | 7.5-22.1 | 13 | <10^-15^ | <0.01 | <0.01 | <0.01 | <0.01 | 0.77 |
| Leaf area (cm^2^) | 481-950 | 656 | <10^-15^ | <0.01 | <0.01 | <0.01 | <0.01 | 0.56 |
| Stem diameter (mm) | 15-30 | 20 | <10^-15^ | <0.01 | <0.01 | <0.01 | <0.01 | 0.58 |
| Crude protein | 2.2-3.5 | 2.5 | <10^-15^ | <0.01 | <0.01 | NS | NS | 0.60 |
| Acid detergent fiber | 9.4-18.8 | 12.5 | <10^-15^ | <0.01 | <0.01 | <0.01 | NS | 0.65 |
| Neutral detergent fiber | 14.5-28.7 | 19.0 | <10^-15^ | <0.01 | <0.01 | <0.01 | NS | 0.69 |

Notes: G$\times$Y means the interaction between genotype and years; G$\times$B indicates the interaction between genotype and block; *h^2^*: broad-sense heritability. NS: Not significant at α = 0.05.

**Supplementary Table 2 The variance of model parameters for sorghum plant height growth**

| **Terms** | **Range** | **Mean** | **Genotype** | **Year** | ***h^2^*** |
| --- | --- | --- | --- | --- | --- |
| *K* | 79–414 | 305 | < 10^-16^ | <10^-13^ | 0.89 |
| *r* | 0.36–0.05 | 0.1 | < 10^-16^ | <10^-6^ | 0.89 |
| *Po* | -1.4– -10.9 | -4.8 | < 10^-16^ | < 10^-16^ | 0.64 |
| Averaged growth rate (cm/day) | 1.6–4.2 | 3.3 | < 10^-16^ | < 10^-16^ | 0.54 |
| Length of high growth rate (days) | 13–54 | 38 | < 10^-16^ | <10^-5^ | 0.79 |
| Max growth rate (cm/day) | 2.8–12.9 | 7.4 | < 10^-16^ | 0.50 | 0.93 |
| Date of max growth rate (Days after planting) | 28–59 | 48 | < 10^-16^ | < 10^-16^ | 0.80 |

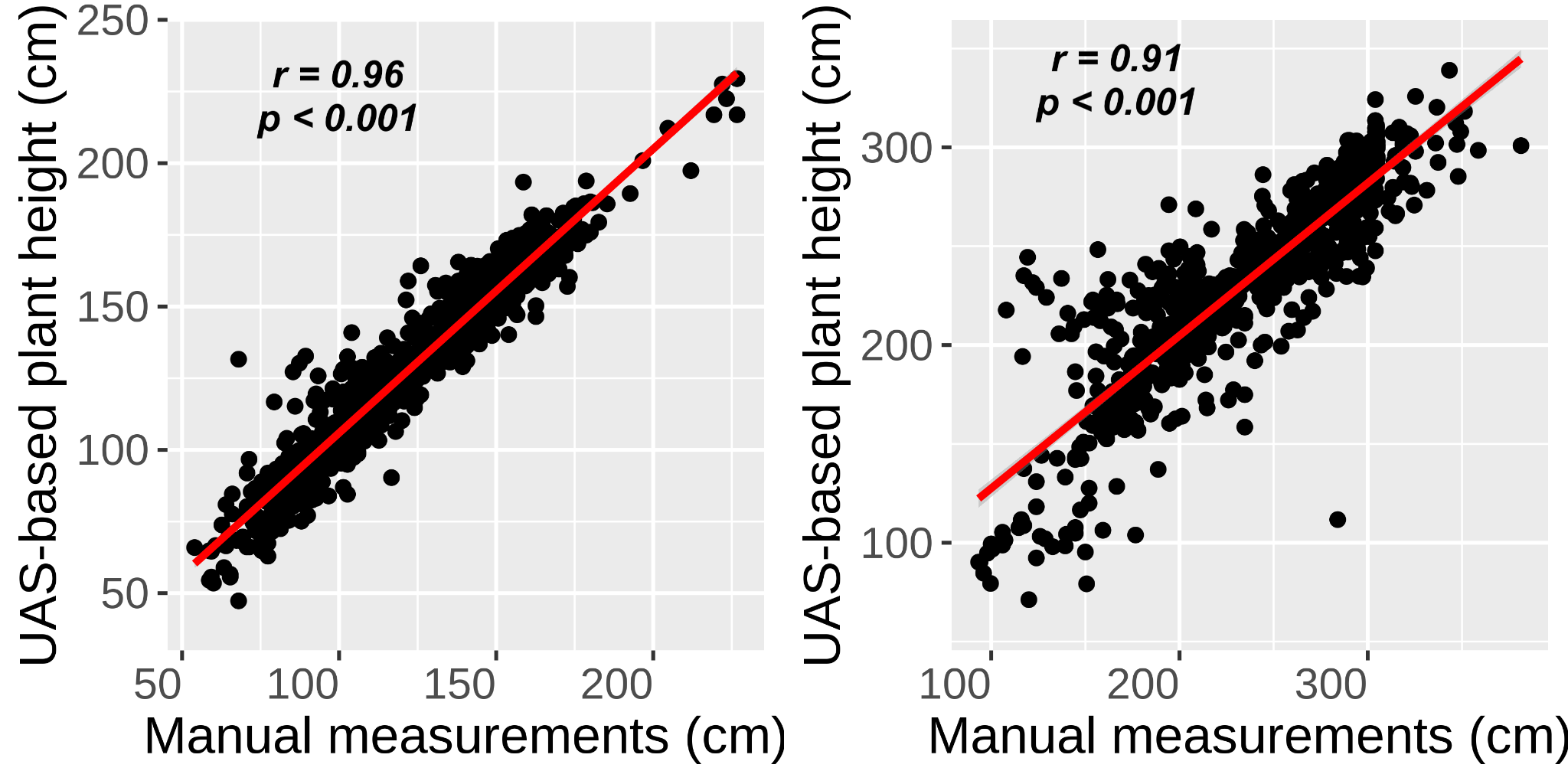

**Supplementary Fig. 1 The correlation between UAS-based plant height and manual measurement at 61 days after planting.**

| 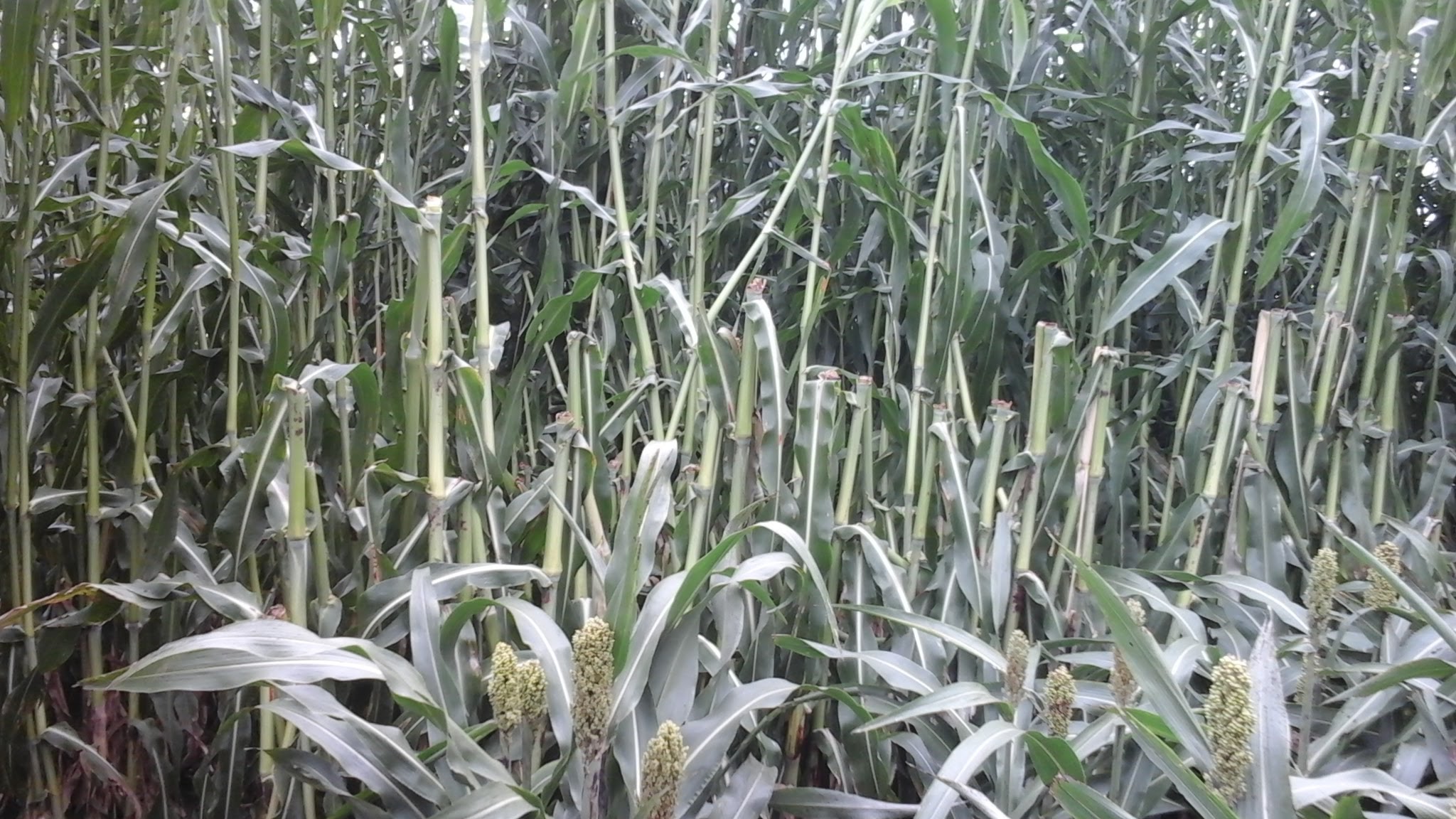 | 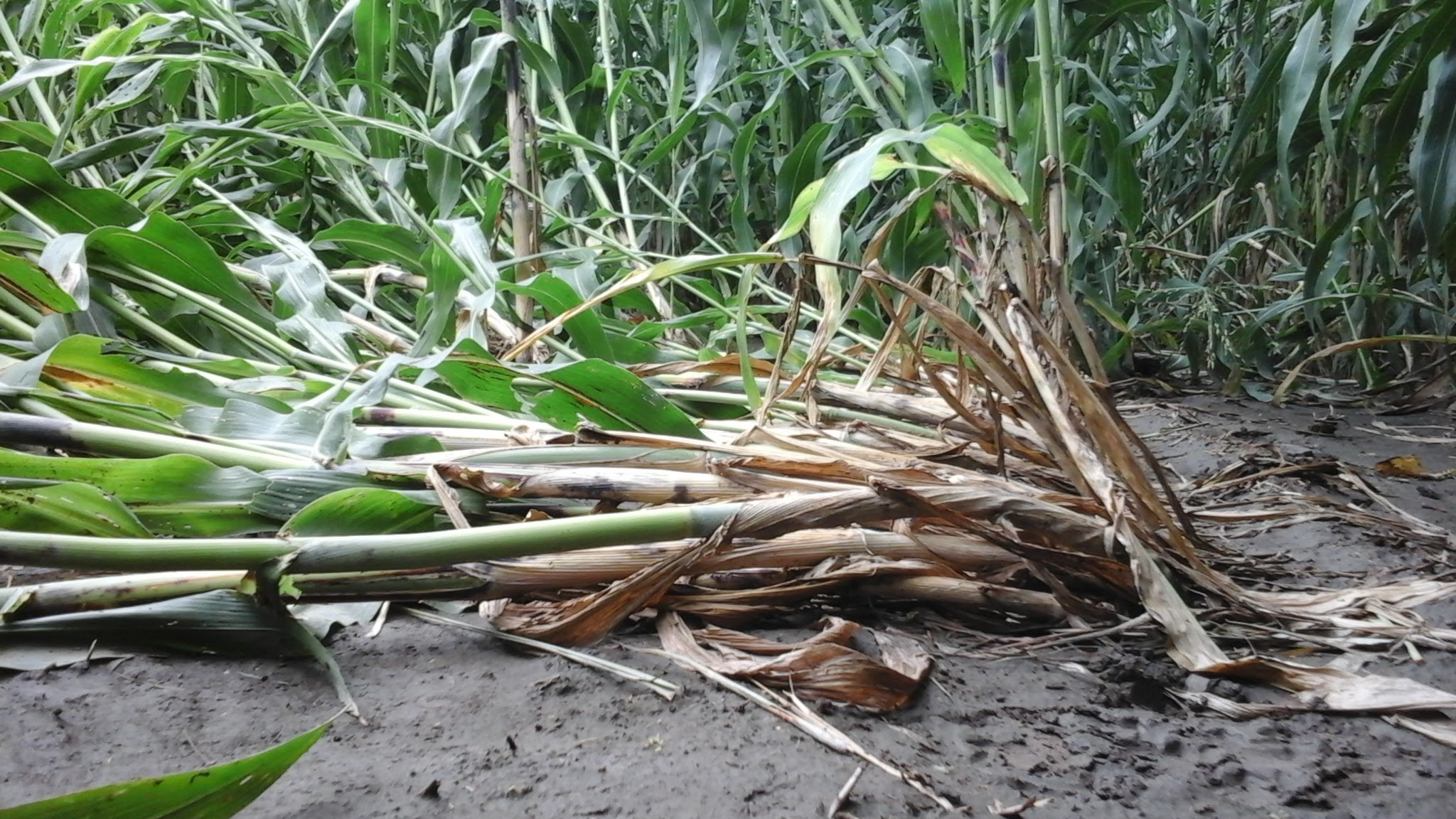 |
| --- | --- |

### **Supplementary Fig. 2 Sorghum lodging in the field.** The left panel is stem lodging and the right panel is shoot lodging.

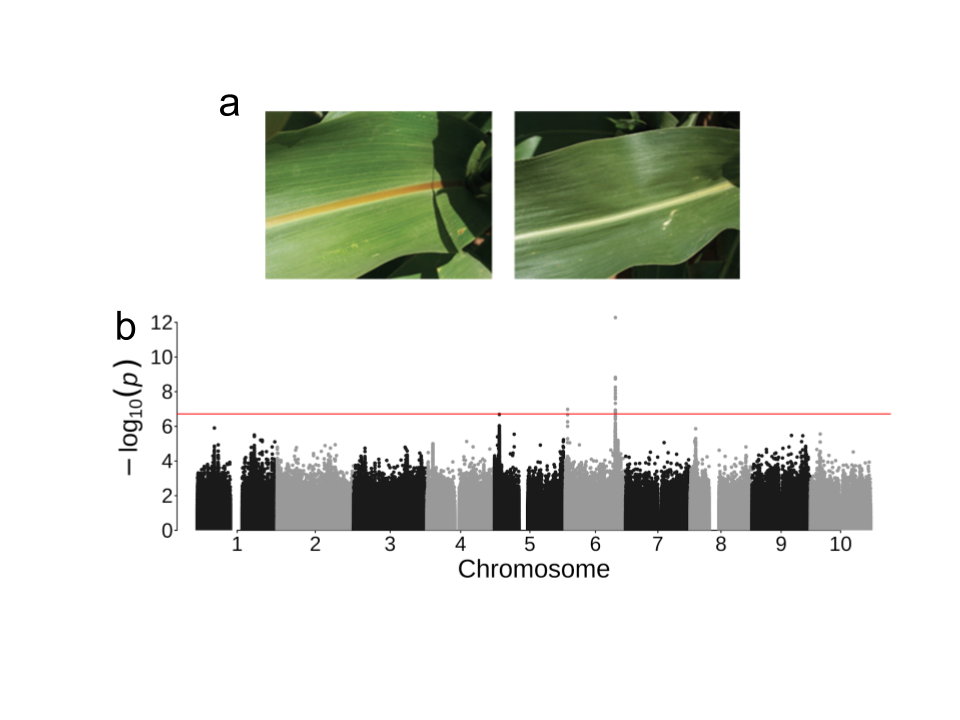

### **Supplementary Fig. 3 Midrib phenotype and genome-wide association study for midrib**. a) The color of the midrib of the leaf is brown (left) and white (right); b) The Manhattan plot of genome-wide association study for midrib using a linear mixed model.

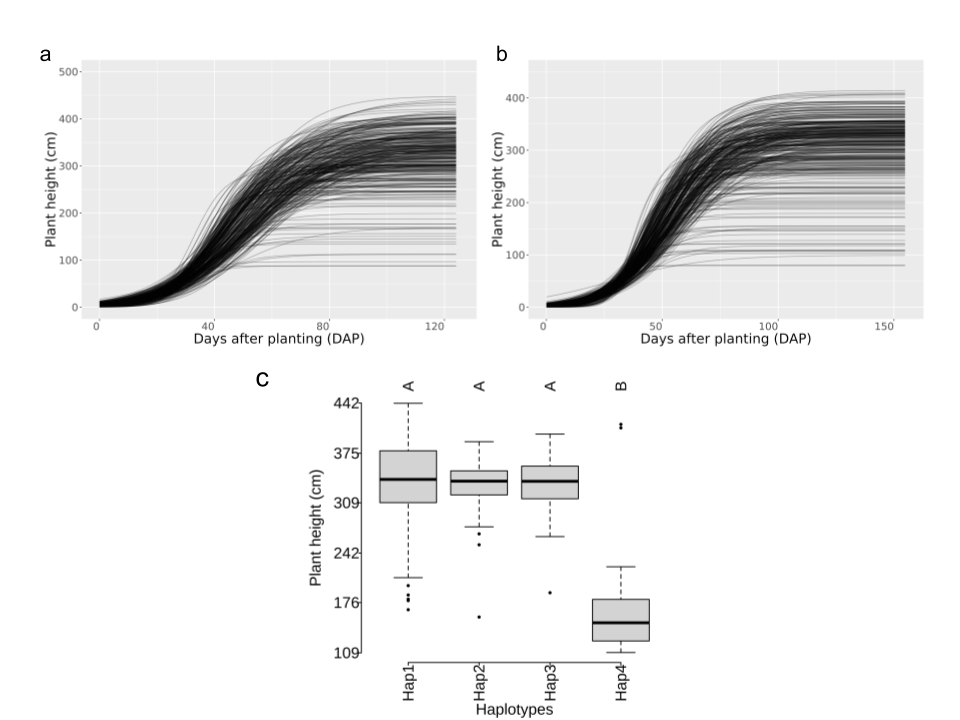

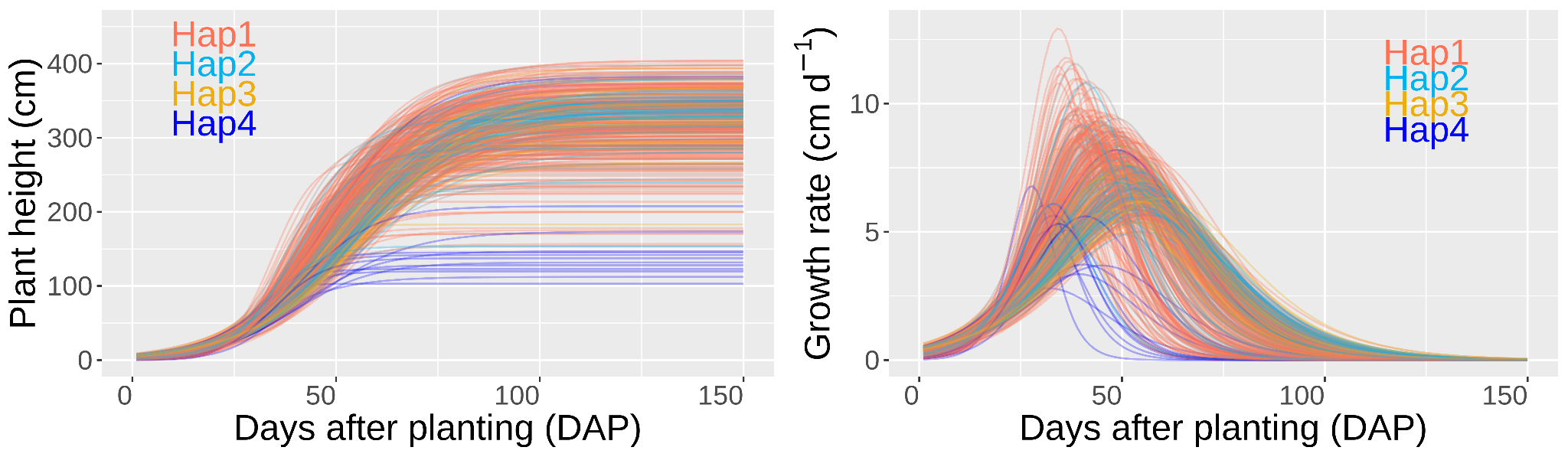

**Supplementary Fig. 4 The growth curve for plant height.** (a) The growth curve for plant height in 2016 and 2017 (b). (c) The averaged modeled growth rate of each accession. The colors indicated four haplotypes of *Dw1*, the major plant height regulator.

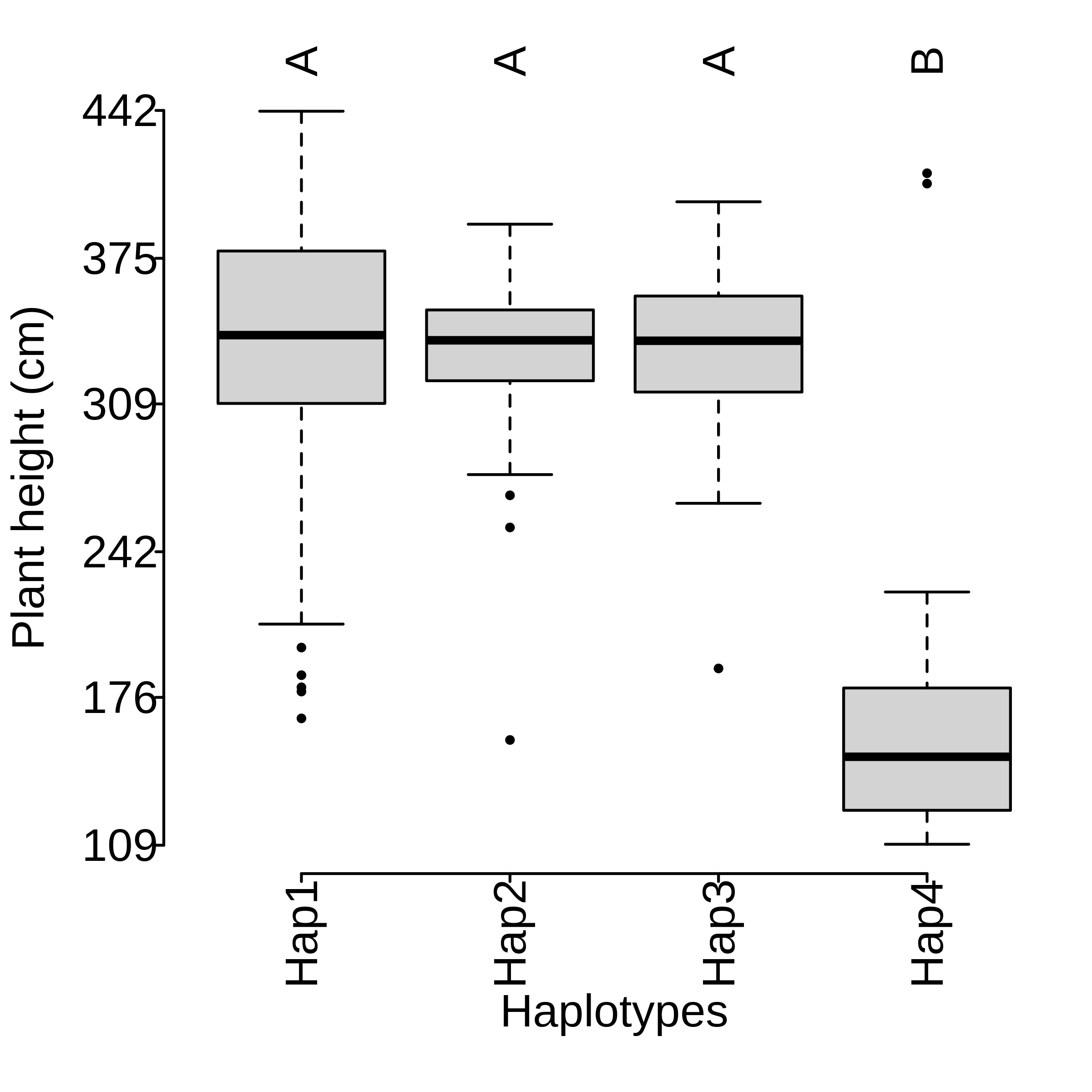

**Supplementary Fig. 5** Plant height comparison among *Dw1* haplotypes*.* The differences among haplotypes were tested using multiple comparisons (Tukey test), and were labeled as 'A', 'B', and 'C'.

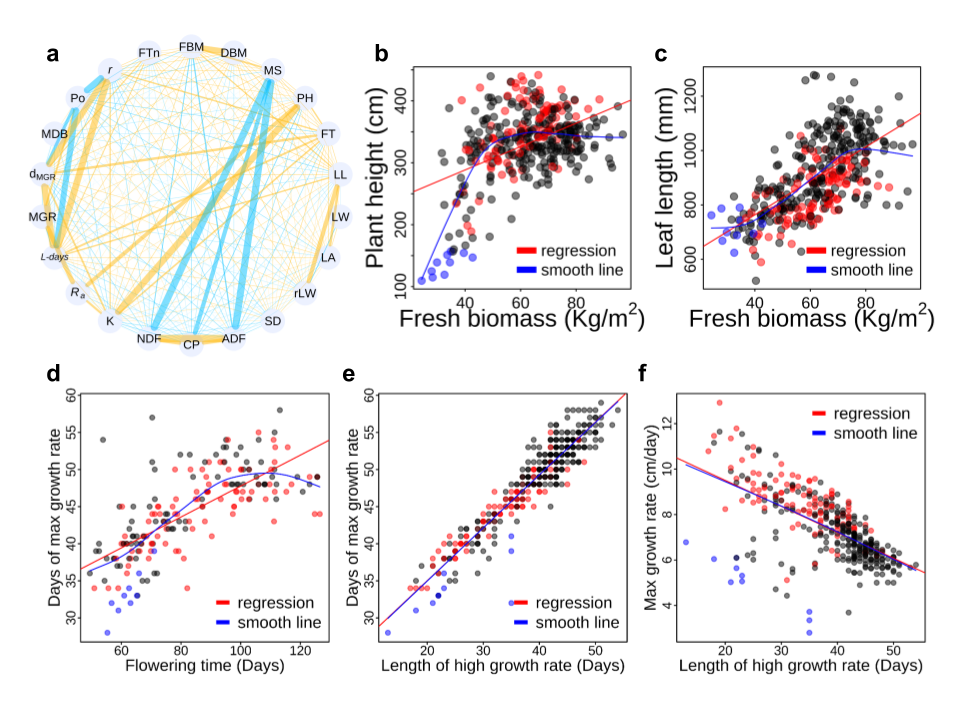

**Supplementary Fig. 6. Trait relationship among yield components, quality components, and dynamic growth parameters underlying yield formation.** (**a**) The correlation network of yield formations. Blue means a negative correlation, yellow indicates a positive correlation and line width is proportional to the correlation. (**b**) The relationship between plant height and fresh biomass. (**c**) The correlation between leaf length and fresh biomass. (**d**) The relationship between the date of MGR and flowering time. (**f**) The relationship between *d_MGR_* and *L-days*. (**e**) The relationship between *L-days* and MGR. The red dots mean sweet sorghum, blue dots mean grain sorghum (defined by < 150 cm). The red line means the regression between two variables, and the blue line means the smooth relation between two variables. Abbreviations: FBM: fresh biomass; DBM: dry biomass; MS: moisture; PH: plant height; FT: flowering time; LL: leaf length; LW: leaf width; LA: leaf area; rLW: ratio of leaf length and leaf width; SD: stem diameter; ADF: acid detergent fiber; CP: crude protein; NDF: Neutral detergent fiber; K: K parameter represented the final plant height; *R_a_*: average growth rate; *L-days*: time length of high growth rate; MGR: max growth rate; *d_MGR_*: the date for max growth rate; MDB: midrib; Po: Po parameter represented the start point of plant height; *r*: *r* parameter represented the intrinsic growth rate of plant height; FTn: flowering or not.

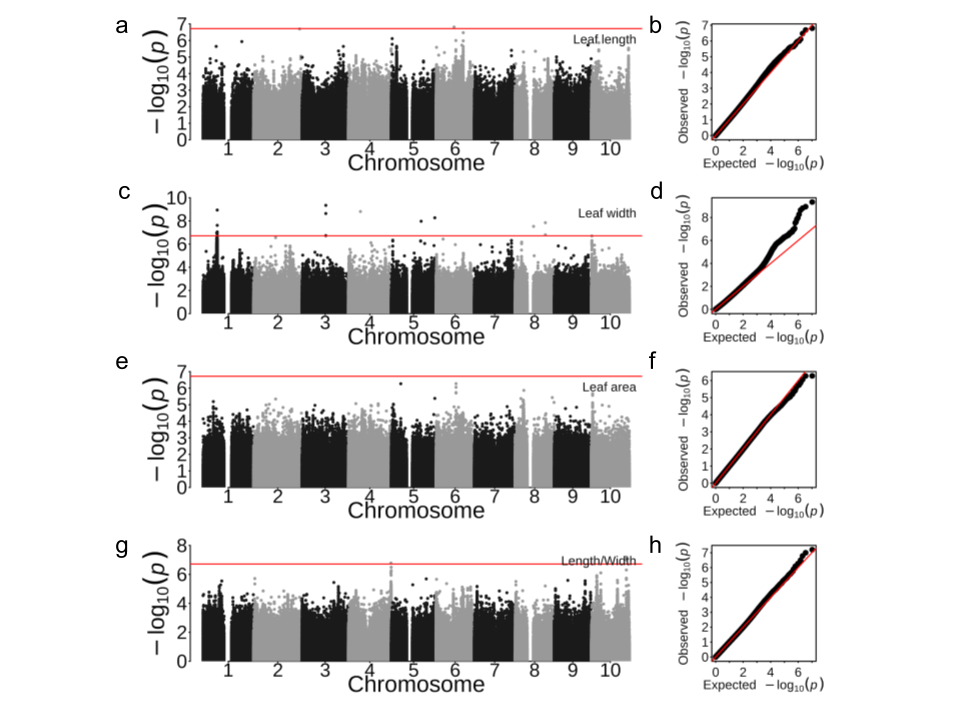

### **Supplementary Fig. 7 Genome-wide association study for leaf traits using the mixed linear model.** a) Manhattan plot for leaf length, b) QQ plot for length width; c) Manhattan plot for leaf area, d) QQ plot for length width; e) Manhattan plot for leaf area, f) QQ plot for leaf area; g) Manhattan plot for the ratio of leaf length to width, and h) QQ plot for the ratio of leaf length to width.

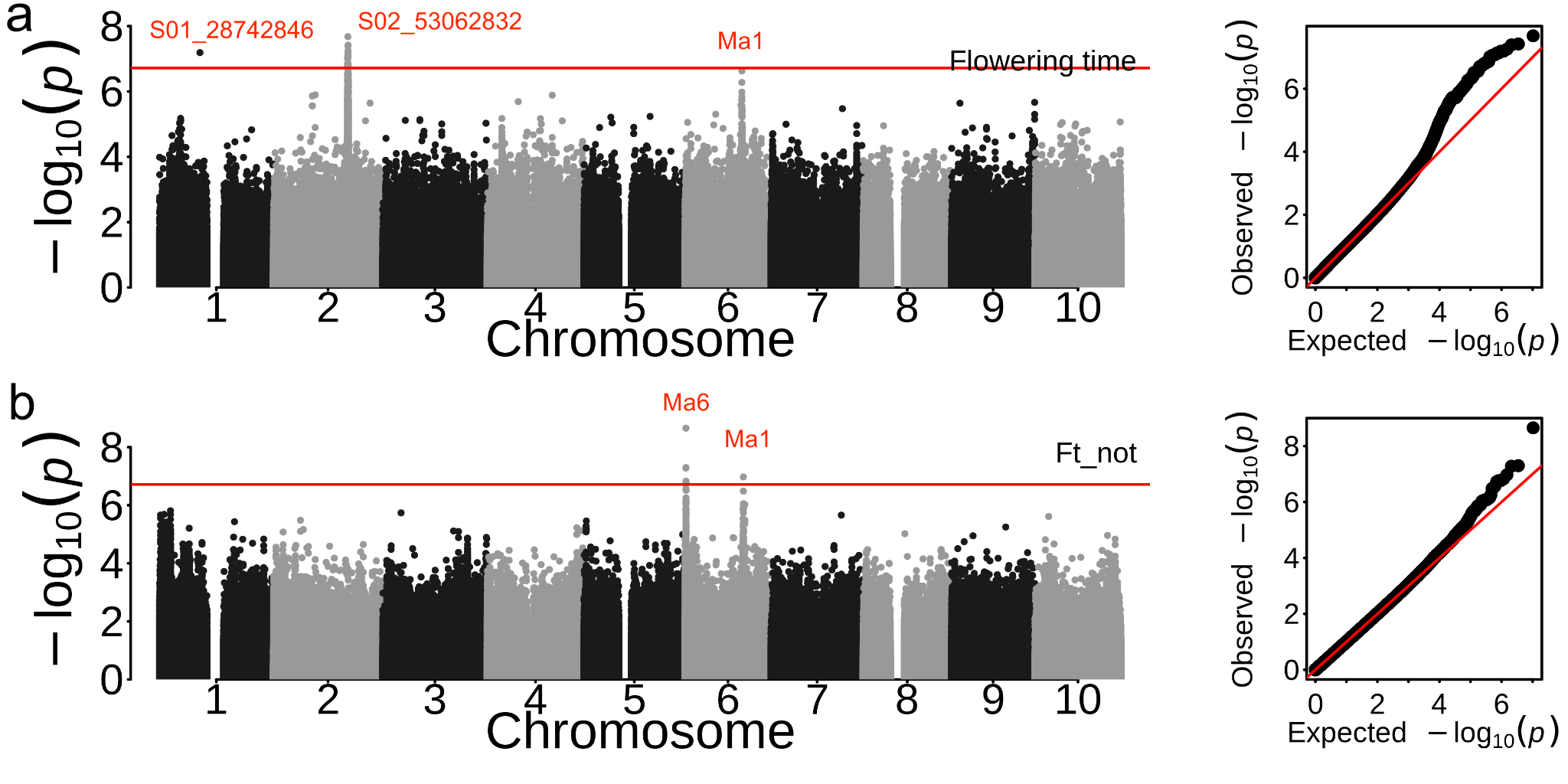

### **Supplementary Fig. 8 Genome-wide association study for flowering time using the mixed linear model.** a) Manhattan and QQ plot of genome-wide association study for flowering time recorded as days after planting. b) Manhattan and QQ plot of genome-wide association study for flowering or not (Ft_not). The known genes were labeled, or leading SNP were labeled.

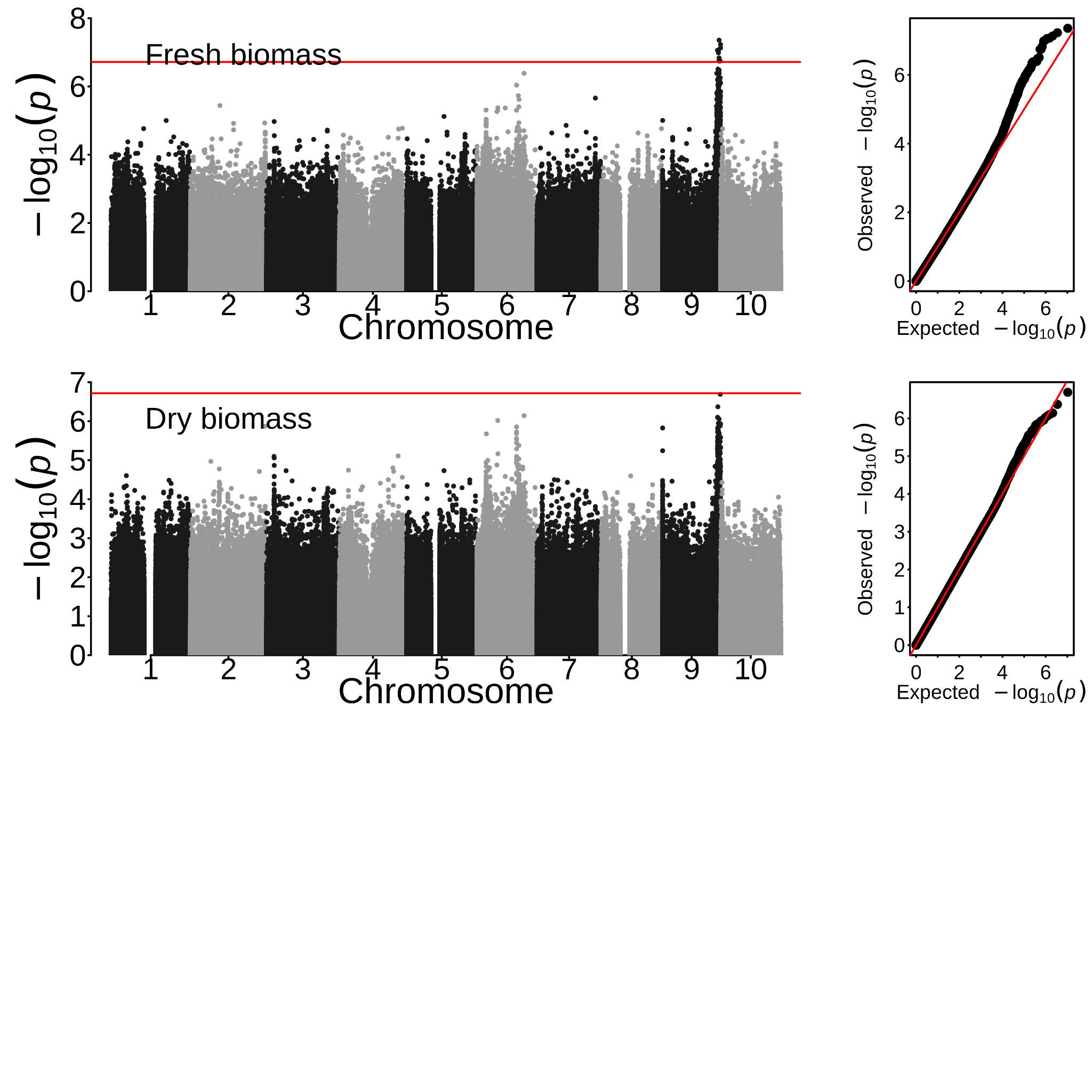

### **Supplementary Fig. 9 The Manhattan plot of genome-wide association study for biomass using the mixed linear model. a) fresh biomass and b) dry biomass.**

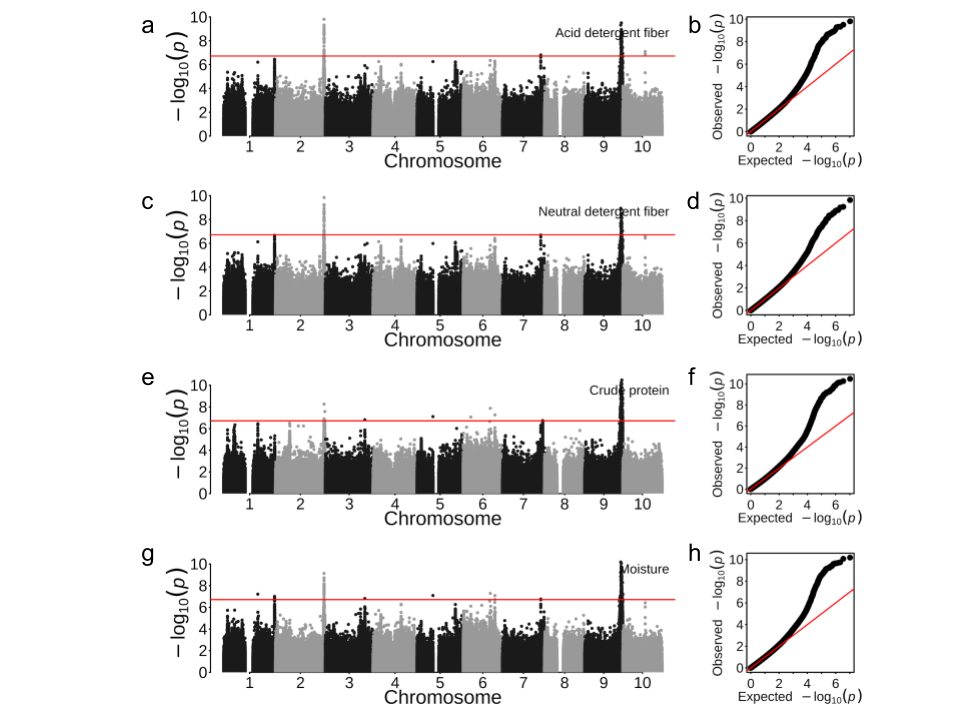

**Supplementary Fig. 10. Genome-wide association study for biochemistry contents of sorghum biomass using the mixed linear model.** a) Manhattan plot for acid detergent fiber, b) QQ plot for acid detergent fiber; c) Manhattan plot for neutral detergent fiber, d) QQ plot for neutral detergent fiber; e) Manhattan plot for crude protein, f) QQ plot for crude protein; g) Manhattan plot for moisture, and h) QQ plot for moisture.

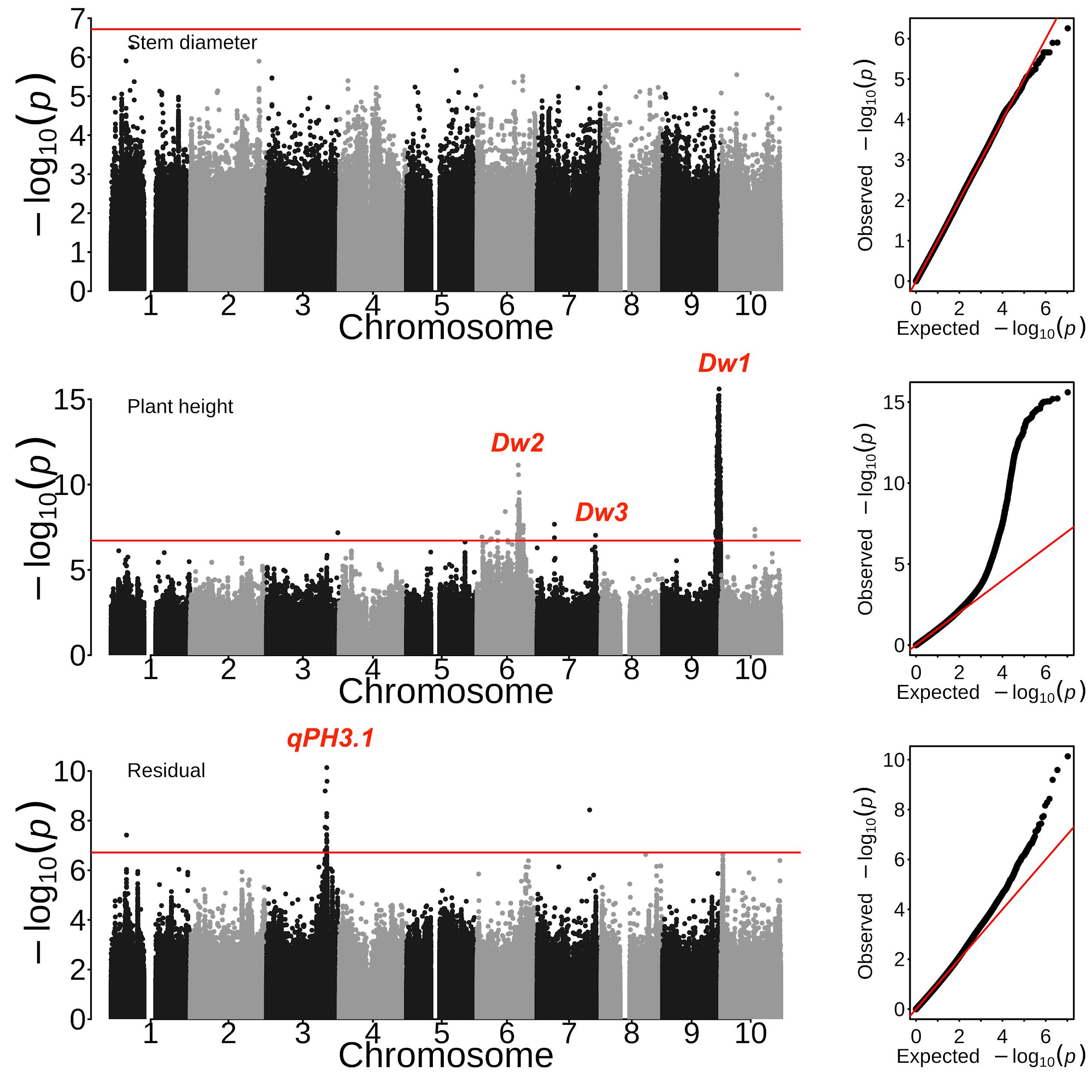

### **Supplementary Fig. 11 Genome-wide association study for stem traits using the mixed linear model.** (a) Manhattan and QQ plot of genome-wide association study for stem diameter, (b) Manhattan and QQ plot of genome-wide association study for plant height.

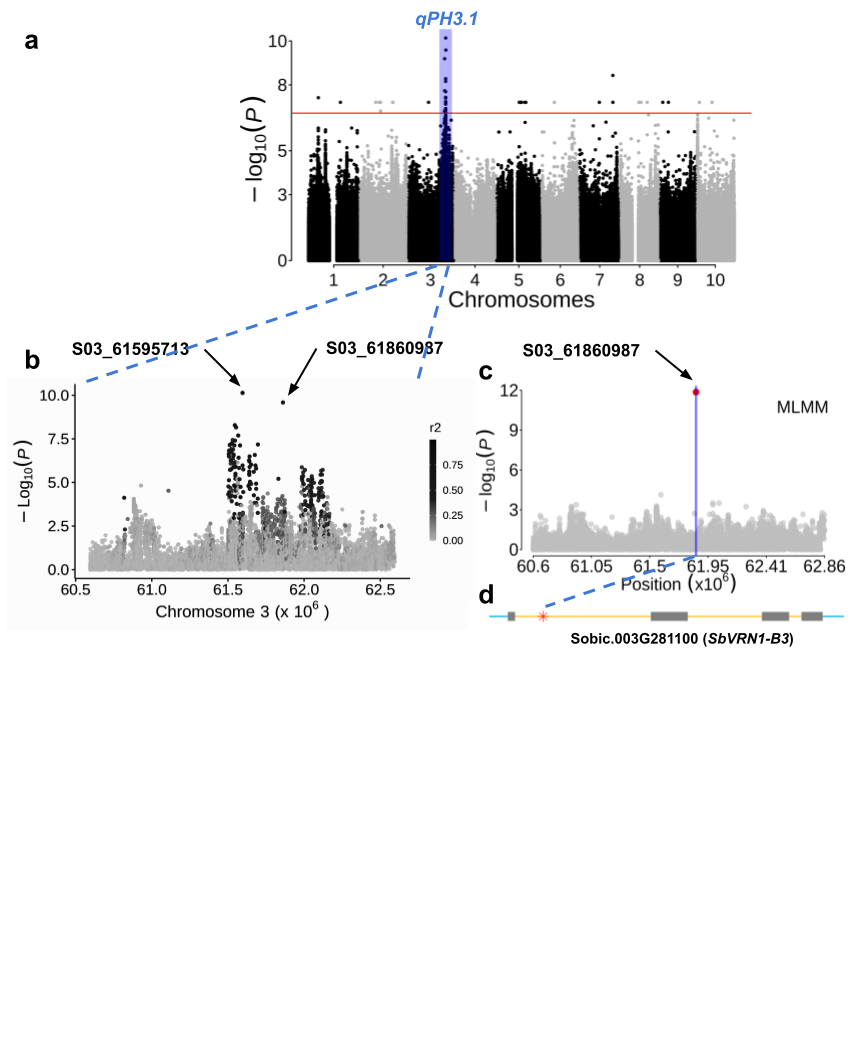

### **Supplementary Fig. 12. GWAS with *Dw1* and *Dw2* as covariates to identify a novel association and candidate gene for plant height.** (**a**) Genome-wide association study of the final manual measurement of plant height using the mixed linear model with *Dw1* and *Dw2* as covariates. The shadow region corresponds to the genomic region by $\pm$1 Mb of leading SNP. (**b**) The zoomed genomic region for association *qPH3.1*. The black-gray color indicated the linkage disequilibrium (*r^2^*) of the given SNP with the leading SNP (S03_61595713). (**c**) The regional association on chromosome 3 for GWAS using a multi-locus mixed model with raw final plant height as phenotype. The leading associated SNP was marked as red (S03_61860987). (**d**) The gene model of the candidate (*SbVRN1-B3*) with the red star indicating the leading SNP S03_61860987. Blue: UTR, yellow: intron, black: exon.

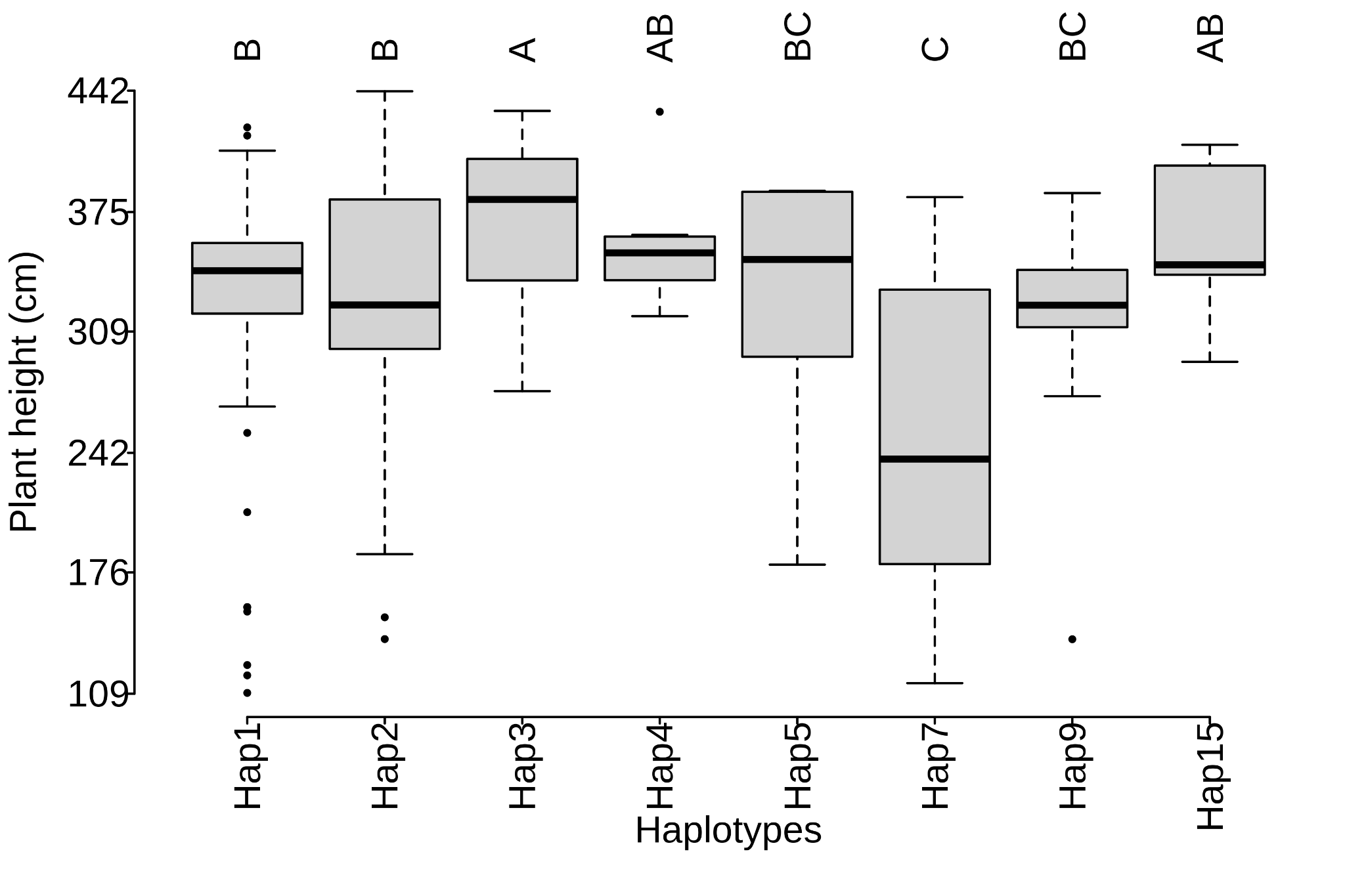

**Supplementary Fig. 13 Plant height comparison among Haplotype of the *qPH3.1* candidate gene (*SbVRN1-B3*; Sobic.003G281100)*.*** The differences were identified using multiple comparisons (HSD.test), and were labeled as 'A', 'B', and 'C'.

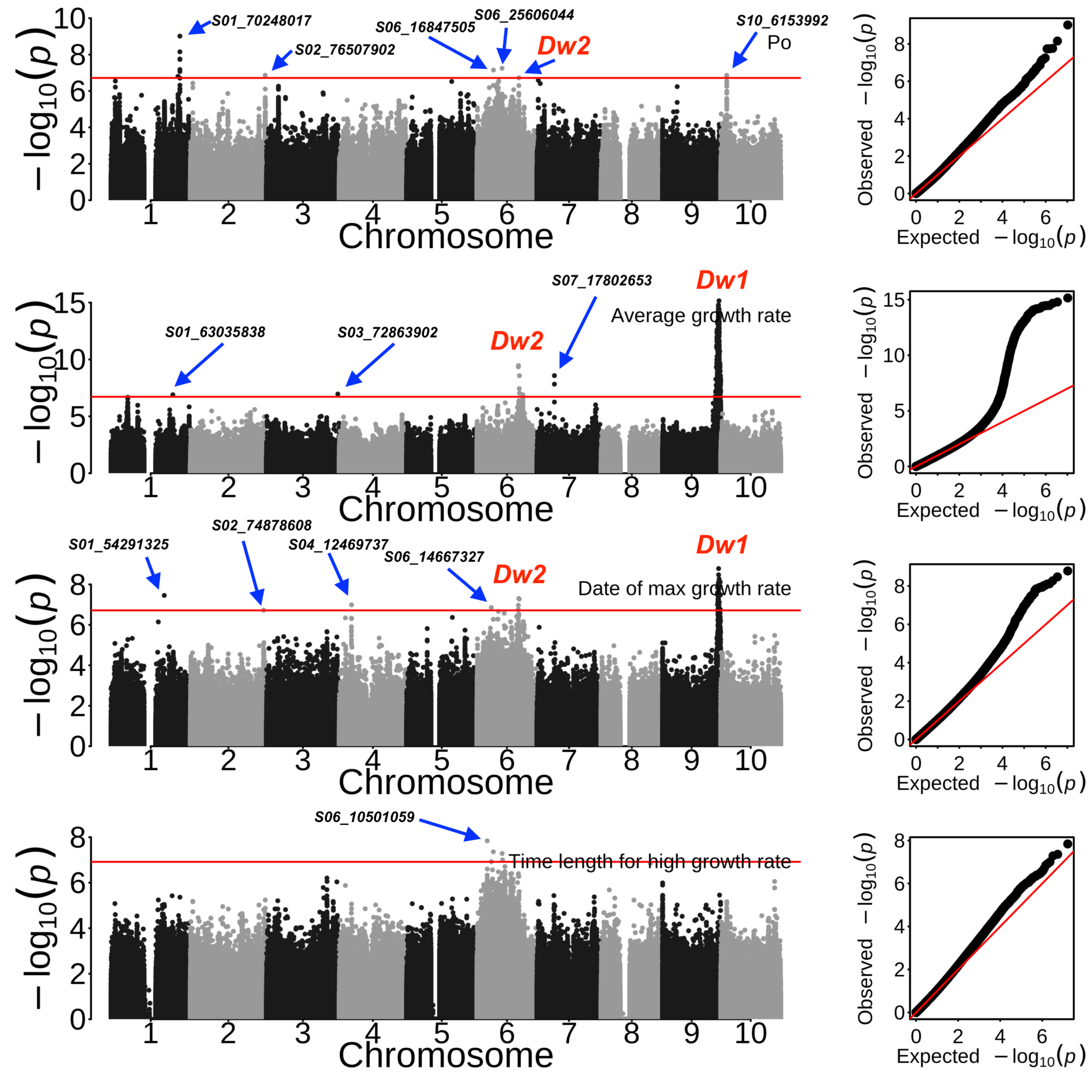

### **Supplementary Fig. 14 Genome-wide association study for plant height growth parameters using the mixed linear model.** a) Manhattan and QQ plot of genome-wide association study for parameter Po, b) Manhattan and QQ plot of genome-wide association study for average growth rate. c) Manhattan and QQ plot of genome-wide association study for data of max growth rate. d) Manhattan and QQ plot of genome-wide association study for time length for high growth rate. The known genes were annotated in red color, the leading SNP were labeled.

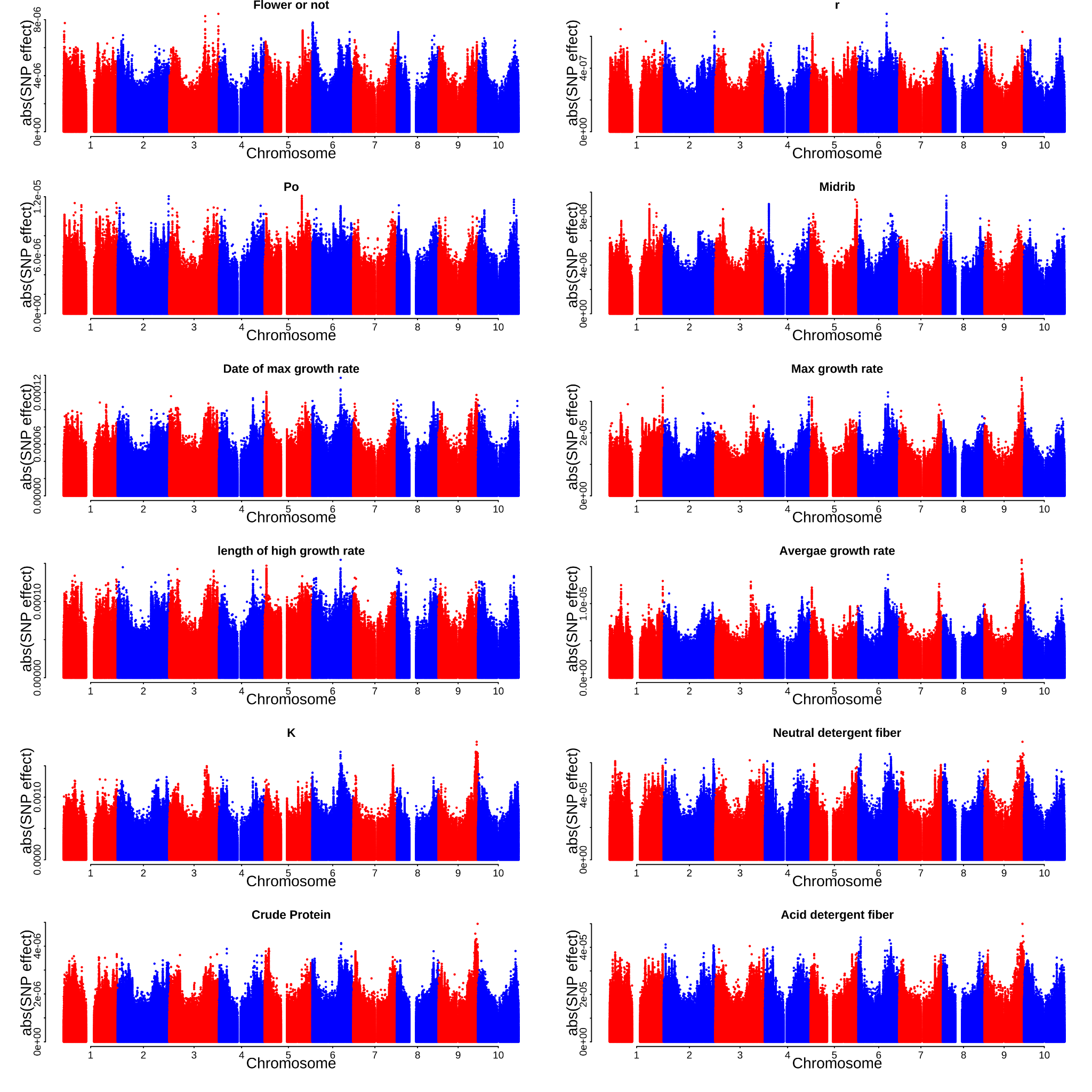

(Continuing)

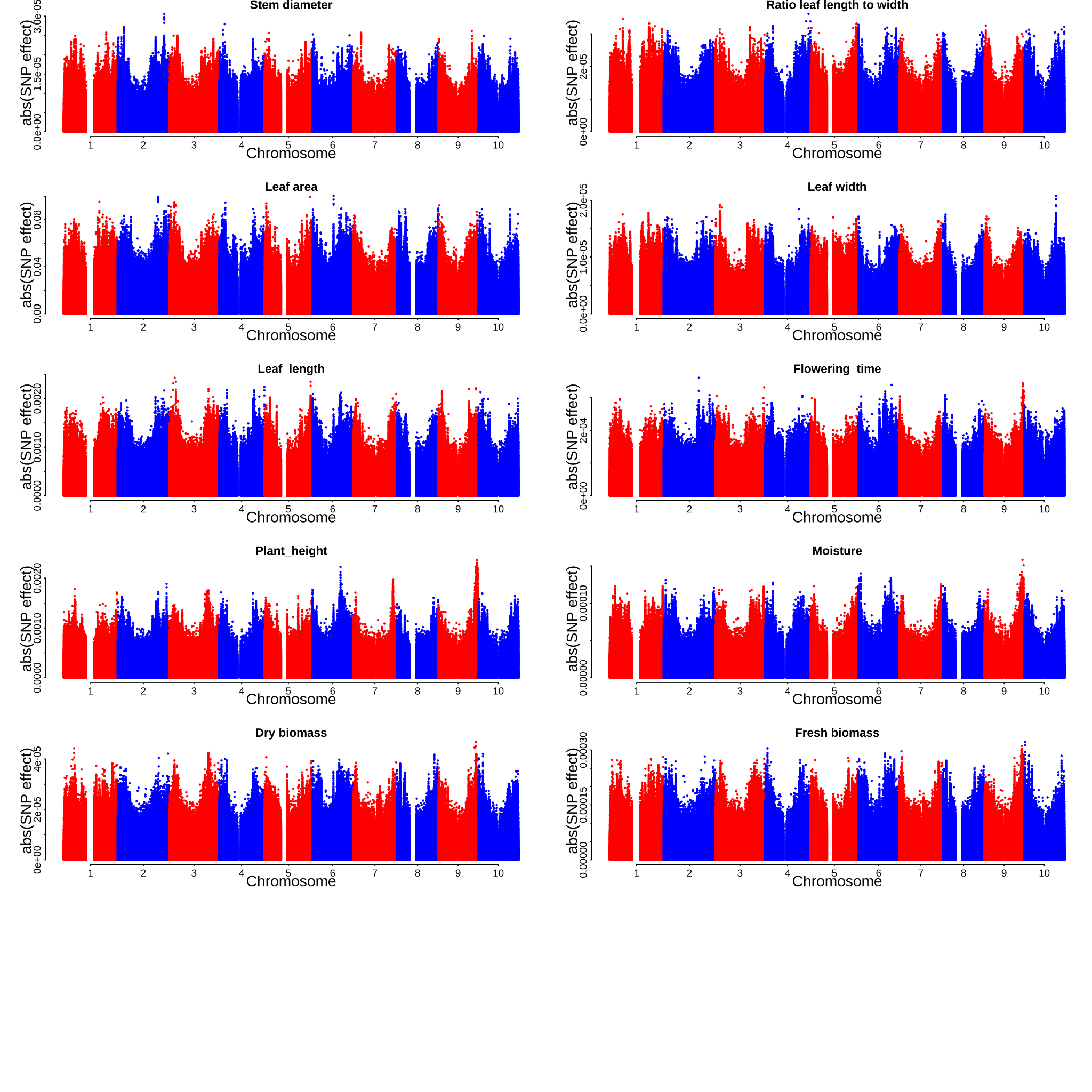

**Supplementary Fig. 15. Genome-wide absolute value marker loadings from GWP models for 22 biomass yield formations in sorghums.** abs(SNP effect) means the absolute value of the marker loading for each SNP.

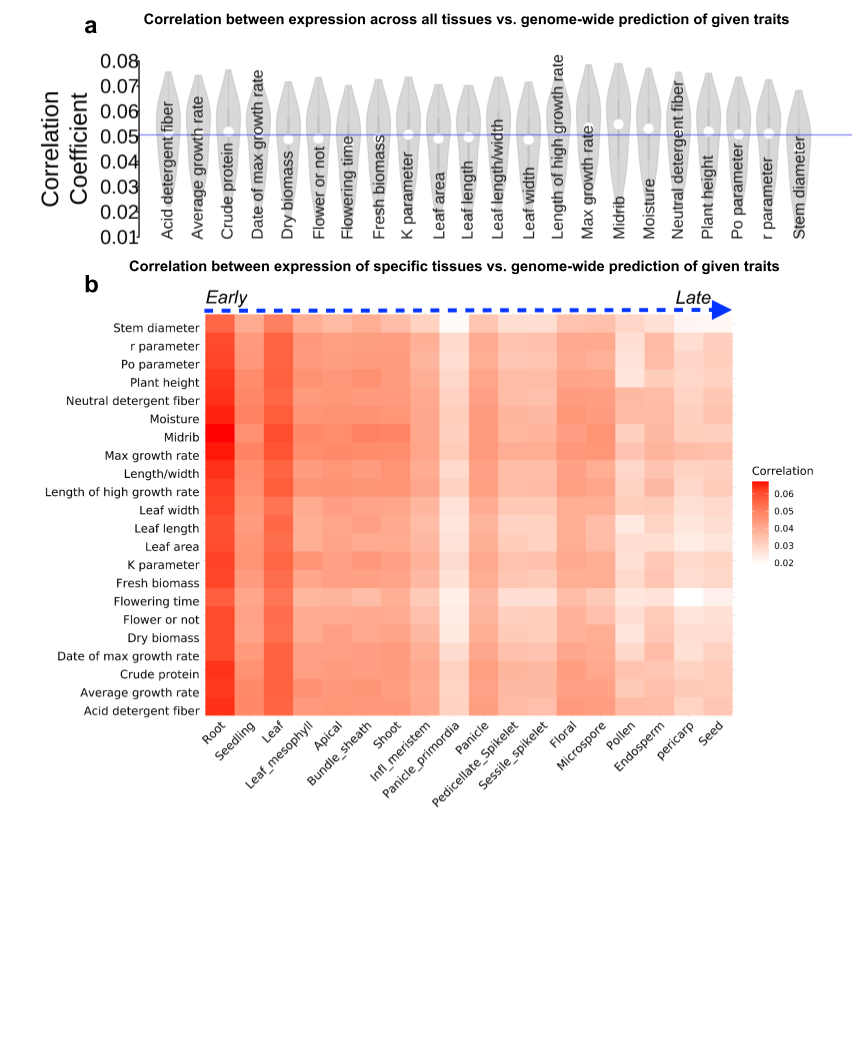

**Supplementary Fig. 16. Testing the omnigenic model based on predicted correlations of polygenic effects and tissue-level gene expression**. The overall correlation coefficient of genome-wide gene expression and gene loadings from a genome-wide prediction. The "gene loading" was calculated using the SNP loading from the genome prediction for SNPs located in a given gene and upstream 5 kb (regulatory region) was used.

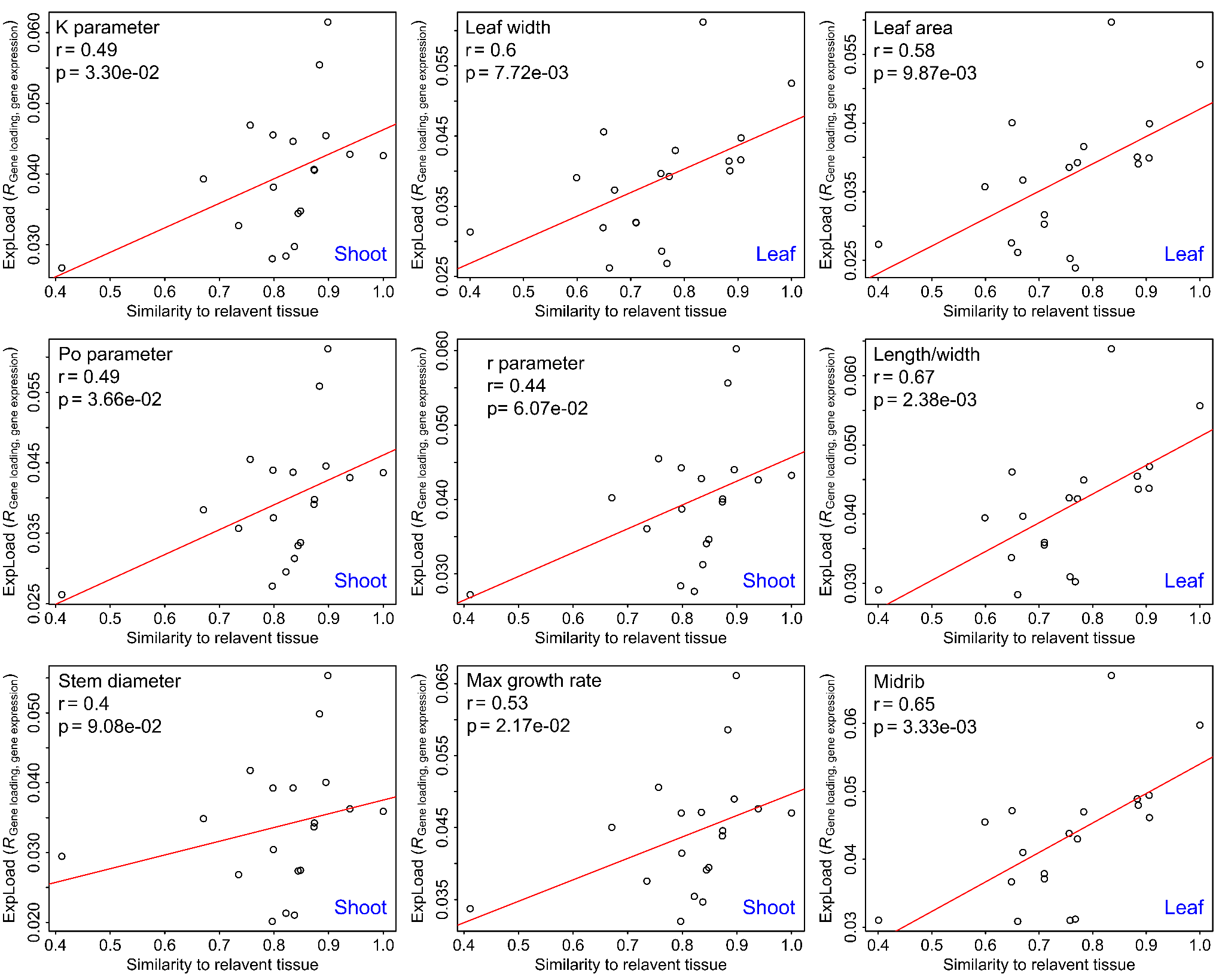

**Supplementary Fig. 17. Putative omnigenic-in-space effects of expression on phenotypic variation.** The correlation coefficient is the correlation between gene loading and expression for traits that easily identify their relevant tissues. The expression space indicates the expression distance between other tissues and the relevant tissue. The relevant tissue was annotated on the top left corner and highlighted as blue text.

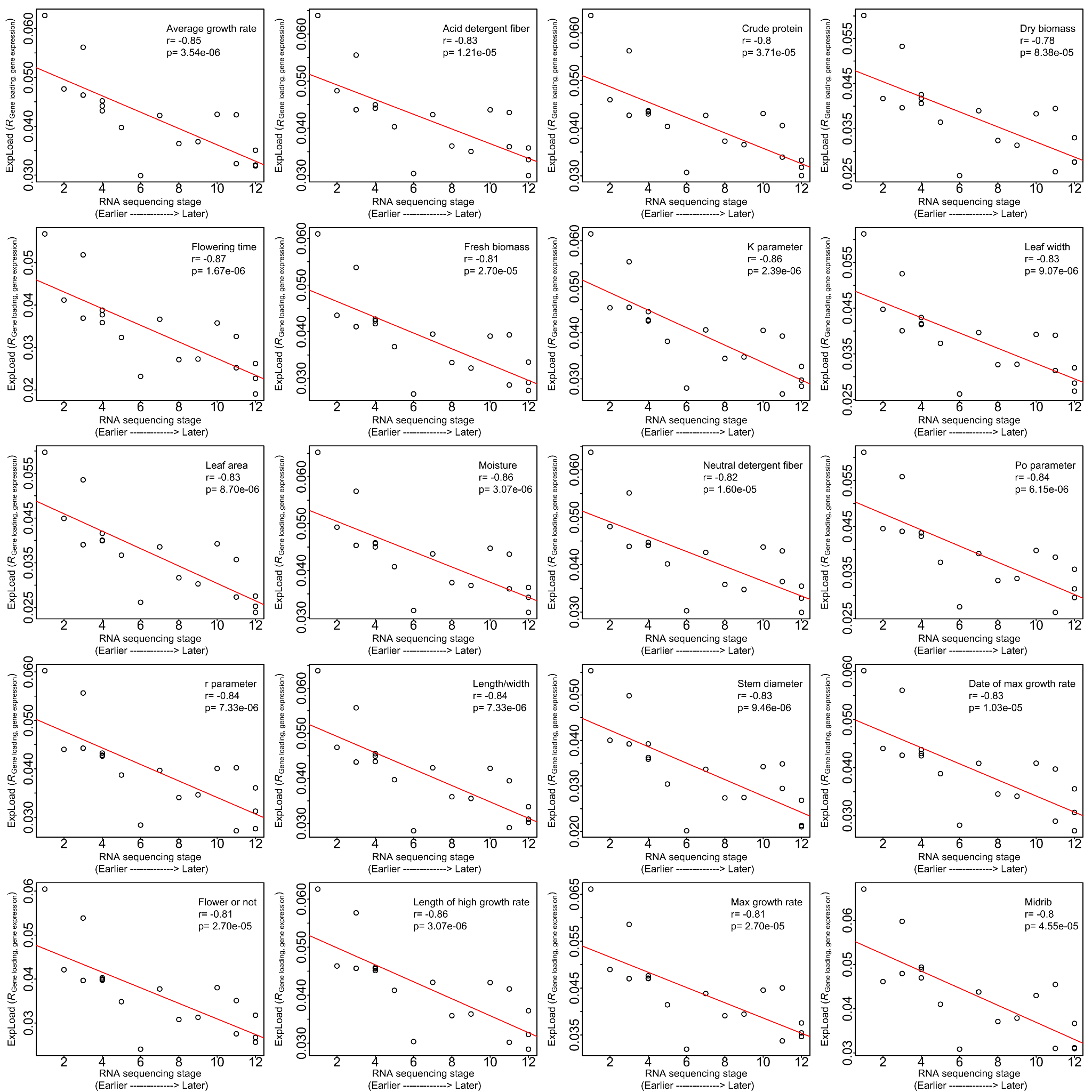

**Supplementary Fig. 18. Putative omnigenic-in-time effects of expression on phenotypic variation.** The corresponding trait for each figure was annotated, the correlation coefficient is the correlation between gene loading and expression. The expression time indicates the order of the tissue generation based on the developmental stage; it ranges from 1 (earlier, root) to 12 (later, seed).
